# Immune activation state modulates the retrieval of infant engrams

**DOI:** 10.1101/2022.04.24.488172

**Authors:** Sarah D. Power, Erika Stewart, Louisa G. Zielke, Eric Patrick Byrne, Clara Ortega-de San Luis, Lydia Lynch, Tomás J. Ryan

## Abstract

Infantile amnesia is possibly the most ubiquitous form of memory loss in mammals. Despite its widespread relevance, little is known about the biological conditions for infantile amnesia to occur and its effect on the engram cells that encode a memory. We investigated how memories are stored in the brain throughout development by integrating engram labeling technology with mouse models of infantile amnesia. Here, we discovered a phenomenon in which male offspring in maternal immune activation models of autism spectrum disorder do not experience infantile amnesia. We rescued the same apparently forgotten infantile memories in mice by optogenetically reactivating dentate gyrus engram cells labeled during complex infant development experiences. Further, we were able to permanently reinstate lost infantile memories by artificially updating the memory engram, demonstrating that infantile amnesia is a reversible process. Our findings suggest that immune activation during development modulates innate, and reversible, forgetting switches that determine whether infantile amnesia will occur.

## Main text

Infantile amnesia is the rapid forgetting of memories formed during early development; and is a largely neglected form of memory loss that seemingly affects 100 % of the human population (*1*, *2*). Not uniquely a human phenomenon, this form of amnesia has been documented in rodents, which show forgetting of contextual and fear memories formed during the infant period (*3–6*). Little is known about the basic neurobiology of infantile amnesia and further its effect on the engrams cell ensembles that encode specific memories. Thanks to the integration of activity-dependent ensemble labeling with optogenetics it is now possible to investigate if a memory engram is still present and/or functional in the brain even in the case of amnesia (*7*). Using this methodology, it has been shown that memory recall can be induced after amnesia by optogenetic activation of engram cells in the hippocampus and other brain regions, demonstrating that these memories are not only still present in the brain,,, but are recoverable (*8–11*). This framework provides an opportunity to investigate how development affects the storage and retrieval of early childhood memories. The environment strongly influences learning and memory (*12*). During development, there are periods in which the developing brain has heightened sensitivity to environmental influences (*6*, *13–18*). Infantile amnesia has been shown to be preventable through pharmacological interventions with GABA agonists (*19–21*) and corticosteroids (*6*) or ectopic administration of neurotrophins (*22*) but the relevance of these switches to organismal function is unknown. Here, we sought to identify naturally occurring variation in infantile amnesia caused by the developmental experience of the animal, and then investigate the effects on engram cell function.

Using a contextual fear conditioning paradigm (Fig. 1B), we trained and tested infant (P17) and adult (P63) mice for fear memory recall 1 or 8 days post-training (Fig. 1C). At both time points, the experimental adult shock group showed significantly more freezing than the no shock control group (Fig. 1C). Infant mice tested for memory recall 1 day following training showed significantly higher levels of freezing compared to the control group (Fig. 1C). While the group tested 8 days after training showed robust infantile amnesia, with similar levels of freezing to the no shock controls (Fig. 1C). Consistent with the literature, infant mice demonstrate forgetting as early as one week after training, while adult mice show continuous memory retention (*4*, *23*).

**Fig. 1.**
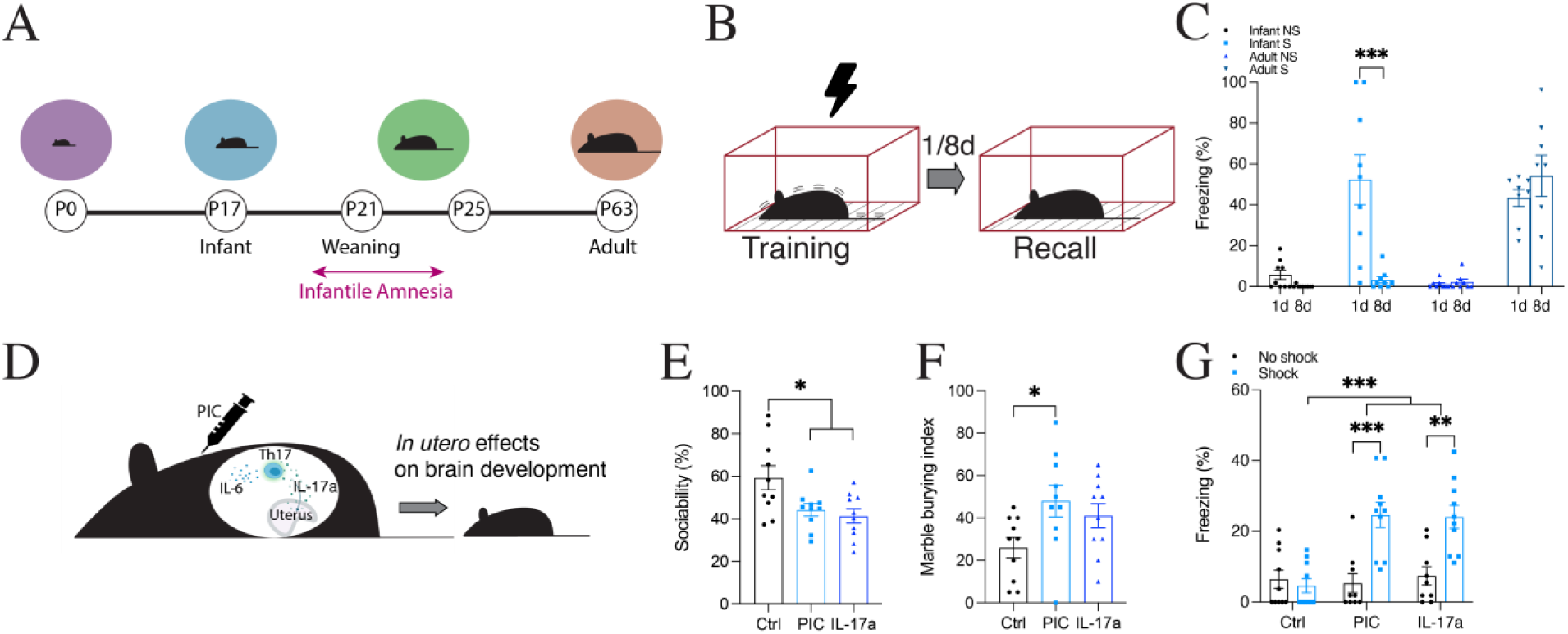
MIA male offspring do not demonstrate infantile amnesia. (**A**) Developmental trajectory of infant mice. (**B**) Behavioral schedule. The black lightning symbol represents foot shocks. Mice are trained using contextual fear conditioning (CFC) in context A and tested for recall 1 or 8 days later. (**C**) Memory recall in context A. Adult (P63) C57BLJ mice (n=8) froze significantly more in context A than no shock controls (n=8) both 1 and 8 days after training. Infant (P17) C57BLJ mice (n=9) froze significantly more than no shock controls during recall 1 day after training. No significant difference in freezing between groups (n=9) when tested for recall 8 days after training. (**D**) Representative diagram of MIA in mice. The syringe indicates Poly I:C (PIC) injection at E12.5. (**E**) Social preference index (% time spent exploring social stimulus out of total object investigation time) of MIA adult male offspring (n=10). (**F**) Marble-burying behavior of MIA adult male offspring (n=10). Percentage of the number of buried marbles is plotted on the y-axis. (**G**) Memory recall in context A after CFC at P17 (n=10). *P < 0.05, P < 0.01, P < 0.001* calculated by one-way (**E**, **F**), or two-way (**C**, **G**) ANOVA with Bonferroni post hoc tests. Data presented as mean ±SEM.

We tested whether altering environmental influences, such as environmental enrichment during infancy and the time of weaning from maternal care, affected the occurrence of infantile amnesia (SFig.1). Since these postnatal interventions had no effect, we next looked at the effect of challenging the embryonic environment. During pregnancy, exposure to pathogens or inflammation during critical periods of brain development (maternal immune activation) has been shown to alter postnatal brain growth and cognitive development (*24*). The viral-mimetic compound Polynosinic-polycytidylic acid (Poly I:C), or the effector cytokine IL-17a, was used to stimulate maternal immune activation (MIA) in pregnant dams at E12.5 in C57BL6/J mice (Fig. 1D). In line with published studies, male, but not female, offspring from pregnant dams treated with Poly I:C or IL-17a demonstrated repetitive behavior and deficits in social behavior, indicative of autism spectrum disorder (ASD) (Fig. 1E, F, SFig. 2A-D) (*13*, *25*, *26*). To test if MIA causes any effect on memory retention, we next trained infant offspring from MIA dams using contextual fear conditioning at P17 (Fig. 1G, SFig. 2E-G). When tested 1 day after training, all cohorts showed similar levels of freezing, irrespective of sex (SFig. 2E, F). However male, but not female offspring from pregnant dams treated with Poly I:C or IL-17a showed retention of infant memory, like that of adult mice, when tested 8 days after training (Fig. 1E, SFig. 2G). This effect was also true when MIA male offspring were tested for recall of a CFC memory 15 days after training (SFig. 2I). Regardless of whether male MIA offspring were tested for recall 8 or 15 days after training, the fear memory was retained. Further, the effect was sex-specific (interaction p = 0.0119) (SFig. J). This represents a novel phenomenon in which male, but not female, offspring from Poly I:C injected dams do not demonstrate infantile amnesia for a contextual fear memory. This phenomenon was also witnessed in offspring from dams injected with IL-17a, indicating the important role of IL-17a signaling.

We characterized and employed an increased engram labeling strategy for whole-brain tagging of engrams cells in infant mice, using a Fostrap transgenic line that expresses ice from an inducible c-fos promoter (Fig. 2A-C) (*27*, *28*). Intraperitoneal (IP) injection of the tamoxifen metabolite, 4-hydroxytamoxifen (4-OHT), allows iCre to translocate into the nucleus. Ai32 transgenic mice express channelrhodopsin-2/EYFP (ChR2-EYFP) following exposure to Cre recombinase (*29*, *30*). Crossing TRAP mice with Ai32 mice makes it possible to produce a stable transgenic line containing both Fos-iCre and ChR2-EYFP. We evaluated the efficiency of the Ai32-FosTRAP engram labeling system in infant (P17) and adult (P42) mice by quantifying the number of ChR2-EYFP^+^ cells in brain regions associated with context and fear memory, including the dentate gyrus (DG), amygdala (AMG), retrosplenial cortex (RSC) and the periaqueductal gray (PAG) at P63 for a home cage, contextual or contextual fear memory (Fig. 2D-F, SFig. 4).

**Fig. 2.**
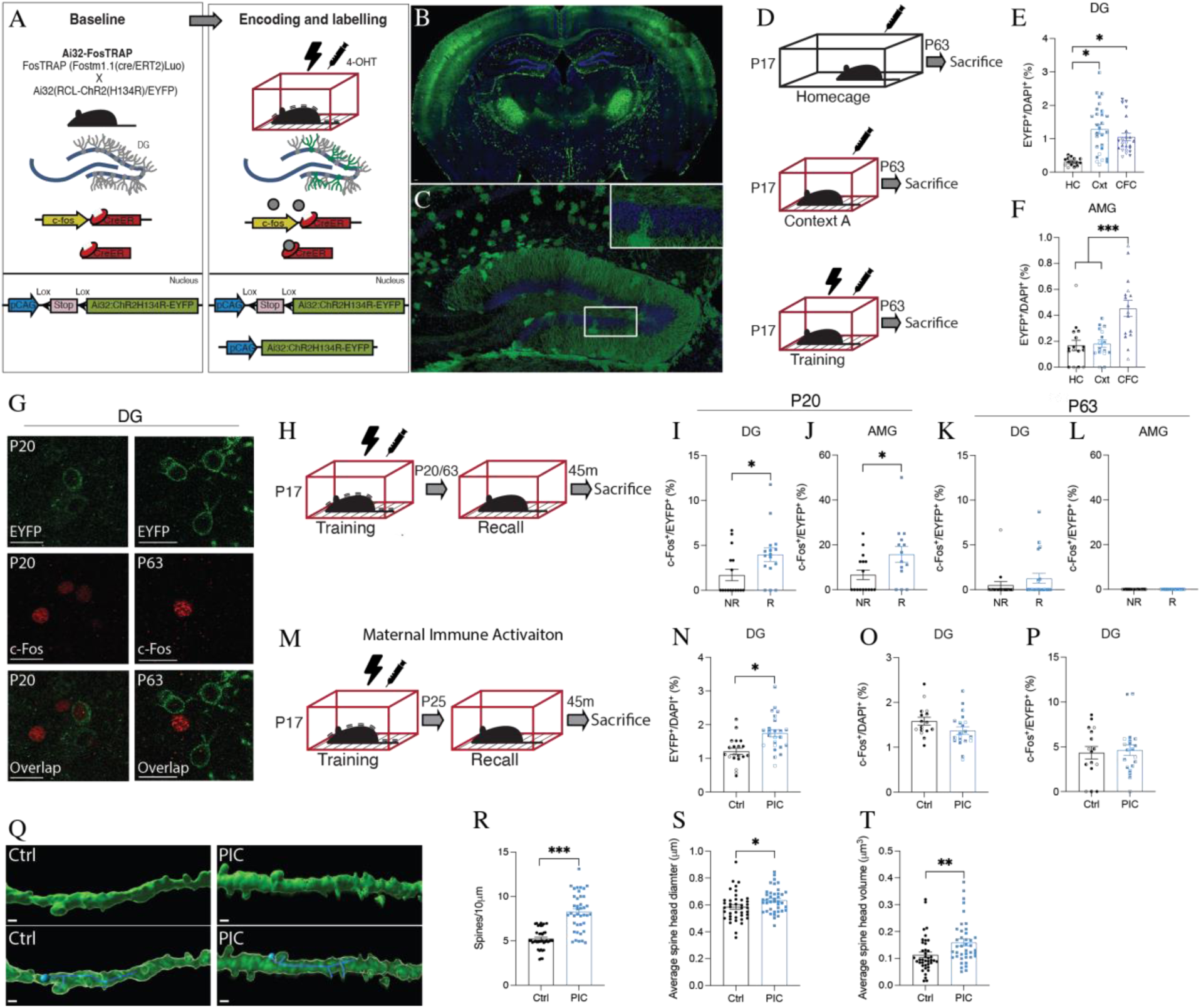
Natural reactivation of engram cells for an infant encoded memory. (**A**) Engram labeling method in Ai32-FosTRAP mice. (**B**) Whole-brain engram labeling of an infant engram. Scale bar 100 μm. (**C**) DG engram cells (ChR2-EYFP^+^ green) in Ai32-FosTRAP mice. Scale bar 100 μm. (**D**) Behavioral schedule for engram labeling of a homecage (HC), contextual (Cxt), or contextual fear memory (CFC) in P17 Ai32-FosTRAP mice. The black lightning symbol represents foot shocks. Syringe symbol represents 4-OHT injection 2 h after training. (**E**, **F**) ChR2-EYFP^+^ cell counts in the DG (N= 4-6, n=4) and AMG (N=4, n=4). (**G**) DG ChR2-EYFP^+^ (green) and c-Fos^+^ (red) cell counts after recall at P20 and P63. Scale bar 20 μm. (**H**, **M**) Behavioral schedule. (**I**, **J**) Engram reactivation in the DG (N=4-6, n=4) and AMG (N=4, n=4) after recall at P20 and P63. (**L**) ChR2-EYFP^+^ cell counts in the DG in MIA male offspring (N=5-6, n=4). (**M**) c-Fos^+^ cell counts in the DG (N=4-5, n=4). (**N**) c-Fos+/ChR2-EYFP^+^ cell counts in the DG (N=4-5, n=4). (**O**) Dendritic spines in MIA male offspring after recall at P25 (N=4, n=10). Scale bar 20 μm (**P**) Spine density per 10μm. (**Q**) Average dendritic spine head diameter (μm). (**R**) Average dendritic spine head volume (μm^2^). *P < 0.05, P < 0.01, P < 0.001* calculated by nested t-test (**E**, **L**-**N**), t-test (**P**-**R**), or nested one-way (**E**, **F**, **I**, **J**) ANOVA with Bonferroni post hoc tests. Data presented as ±SEM.

We next investigated whether engram cells formed at the time of infant memory encoding are reactivated by exposure to training cues (Fig. 2H). Cellular activation of engram cells was examined by quantifying the number of EYFP^+^ c-Fos^+^ cells before (P20) and after (P63) the infantile amnesia period across multiple brain regions (Fig. 2G, SFig. 5-6). Natural recall cues result in above chance EYFP^+^/c-Fos^+^ overlap in the DG and AMG at P20 (Fig. 2I, J, SFig. 5C, F), an effect that was not seen at P63 (Fig. 2I, J, SFig. 6C, F). We next compared engram reactivation after natural recall for a contextual fear memory in male offspring from MIA dams (Fig. 2K). Interestingly, male, but not female, offspring from MIA dams have an increased number of EYFP^+^ engram cells in the DG compared to controls (Fig. 2L, SFig. 2H), indicating a larger engram ensemble also reported for a genetic model of ASD (*31*). However, no difference was seen in the relative level of engram reactivation between groups as both demonstrated an above chance level of EYFP^+^/c-Fos^+^ overlap in the DG (Fig. 2N). Dendritic spine analysis indicated that engram cells in MIA offspring had significantly increased dendritic spine density (Fig. 2P) as well as larger spine head diameter and volume relative to control animals at P25 (Fig. 2Q, R). Thus, a combination of larger engram size, and increased dendritic plasticity, may contribute to functional engram expression of infant engrams in MIA offspring.

To investigate the functionality of memories after naturally occurring infantile amnesia, we tested the behavioral effect of optogenetically stimulating infant-labeled DG engram cells in adults. Engram cells for context A were labeled in Ai32-FosTRAP mice at P17 using CFC (Fig. 3A, B). Although both cohorts were placed in context A, only the experimental group received foot shocks (S). At P63, both cohorts were placed in context C for a 12 min test session where they received four 3 min epochs of blue light on or off. During the light-off epochs, both groups showed background levels of freezing (Fig. 3D). The experimental group showed significantly higher levels of freezing during the light-on epochs (Fig. 3D, E). Despite the same number of EYFP^+^ cells being labeled (Fig. 3F), this light-induced freezing was not observed for animals that underwent the same behavioral paradigm but did not receive any footshocks at P17. While stimulation of a neutral memory, labeled in infancy, does not result in freezing behavior (SFig. 9A-C). We extended this paradigm by optogenetically stimulating infant-labeled CA1 engram cells (SFig. 9D). Optogenetic stimulation at 4 Hz (SFig. 9H, I), but not 20 Hz (SFig. 9F), of infant-labeled engram cells in the CA1 of adult mice resulted in light-induced freezing. Based on these data, ‘lost’ infantile memories can be acutely recalled by optogenetic stimulation in either the DG or CA1.

**Fig. 3.**
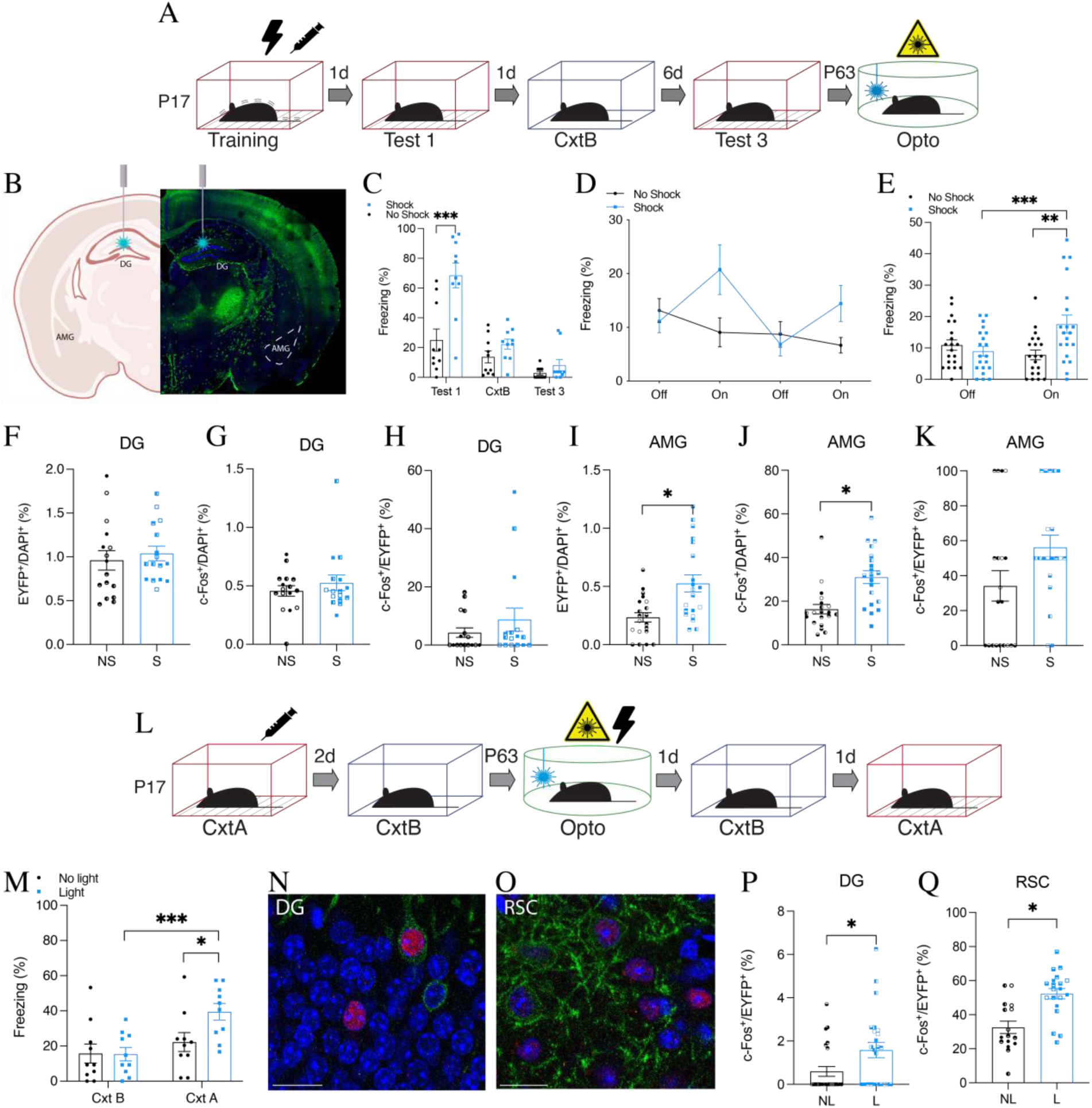
Artificial updating of infant encoded engram cells permanently reinstates a ‘lost’ infantile memory. (**A**) Behavioral schedule for optogenetic reactivation of infant DG engram cells in adulthood. (**B**) Representative image of optogenetic stimulation of DG engram cells in Ai32-FosTRAP labeled mice. (**C**) Freezing levels of infant mice (n=10) during natural memory recall. Shock (S), no shock (NS). (**D**) Memory recall in context C (engram reactivation) with light-off and light-on epochs. (**E**) Freezing for the two light-off and light-on epochs averaged. (**F**, **I**) ChR2-EYFP^+^ cell counts in the DG (N=4, n=4) and AMG (N=5, n=4) after optogenetic stimulation at P63. (**G**, **J**) c-Fos^+^ cell counts in the DG and AMG. (**H**, **K**) c-Fos^+^/ChR2-EYFP^+^ cell counts in the DG and AMG. (**l**) Behavioral schedule for artificial updating of an infant engram. (**M**) Freezing levels during recall in context B and context A (n=10). Experimental light group froze significantly more in context A. (**N**, **O**) Histological representation of engram reactivation in the DG (**N**) and RSC (**O**) during recall in context A. DAPI^+^ (blue), ChR2-EYFP^+^ (green), c-Fos^+^ (red). Scale bar 20 μm. **(P, Q**) DG (N=6, n=4) and RSC (N=4-5, n=4) c-Fos^+^/ChR2-EYFP^+^ cell counts after recall in context A. *P < 0.05, P < 0.01, P < 0.001* calculated by nested t-test (**F**-**K**, **P**, **Q**) or two-way ANOVA (**C**, **E**, **M**) ANOVA with Bonferroni post hoc tests. Data presented as ±SEM.

Although optogenetic stimulation of a forgotten infant memory in adulthood resulted in a specific behavioral response, it is unclear how the information survives. We investigated the connectivity between engram cells in downstream regions, after infantile amnesia, by histologically assessing engram cell reactivation following optogenetic stimulation of upstream engram cells (Fig. 3F-K) (*8*, *32*). The resulting c-Fos^+^ counts were equivalent in the hippocampus after light activation of a CFC (shock, S) or a contextual (no shock, NS) memory (Fig. 3G). In contrast, c-Fos^+^ cell counts were significantly higher in the AMG after light activation of an infant CFC memory (Fig. 3J), demonstrating that amygdala activity correlates with the behavioral expression of freezing. Light activation of DG engram cells results in an above chance c-Fos^+^/EYFP^+^ overlap in the DG (Fig. 3H, SFig. 7B) and crucially in other downstream brain regions including the AMG (Fig. 3K, SFig. 7C), RSC (SFig. 7F, G) and PAG (SFig. 7J, K). These results demonstrate that the connectivity pattern of the engram survives infantile amnesia and persists into adulthood, and optogenetic stimulation of a lost infantile memory is sufficient to reactivate the functional connectivity patterns. Further, optogenetic stimulation of an engram encoded after (P29) the infantile amnesia period also increased engram reactivation in the DG, AMG, RSC, and PAG (SFig. 8A-Q).

Since optogenetic stimulation results in memory recall under artificial conditions, we sought to reinstate an infant memory permanently by adopting an optogenetic induction procedure to induce plasticity in engram cells (SFig. 10A, B) (*8*, *33*). Infant mice can form purely contextual memories that can be updated with shock information (SFig. 10C-F). At P63 and in a novel context, we optogenetically stimulated DG engram cells for context A, originally encoded at P17, while simultaneously delivering footshocks to create a false association of an infant context engram with adult shock experience (Fig. 3L). Animals subsequently froze significantly more when exposed to context A even though they were shocked in context C (Fig. 3M). This increased freezing was not due to generalization, since the control group did not freeze in context B. There was also an increased level of overlap of c-Fos^+^/EYFP^+^ cells in the DG and RSC after exposure to context A, demonstrating that permanent engram reinstatement was also reflected at a cellular level (Fig. 3N-Q, SFig. 11A-N). This histological result demonstrates that engram cells active at the time of encoding were reactivated during adult exposure to context A. Together, these findings show that the plasticity that accompanies the targeted updating of an infant engram restores the natural accessibility of that engram to the appropriate perceptual cues, and that context specificity is maintained into adulthood.

Although fear conditioning is a robust phenomenon, we wanted to further our knowledge by testing the relevance of infantile amnesia to other types of memory. To do this, we next looked at this form of forgetting in a novel object recognition behavioral paradigm (Fig. 4A). Although studies have looked at novel object location in infant rats, there has been no clear demonstration of infantile amnesia for novel object recognition in mice at this stage of development (*34*, *35*). Infant mice demonstrated rapid forgetting for novel object recognition tasks, following the developmental trend like that of other forms of episodic memory. Both adult and infant cohorts spent more time exploring a novel object relative to a familiar object when tested 1 day after acquisition, demonstrating that a long-term memory was formed (Fig. 4B, SFig. 12A-C). When tested again 8 days later, only the adult cohort spent significantly more time exploring the novel object (Fig. 4B, SFig. 12C). We then investigated whether male offspring from MIA dams show infantile amnesia for object memory (Fig. 4C). When tested for object memory retention, male offspring from MIA dams spent significantly more time with the novel object 8 days after acquisition (Fig. 4D, SFig. 12E). Like contextual fear memory, male offspring from Poly I:C or IL-17a injected dams do not show infantile amnesia for an object memory.

**Fig. 4.**
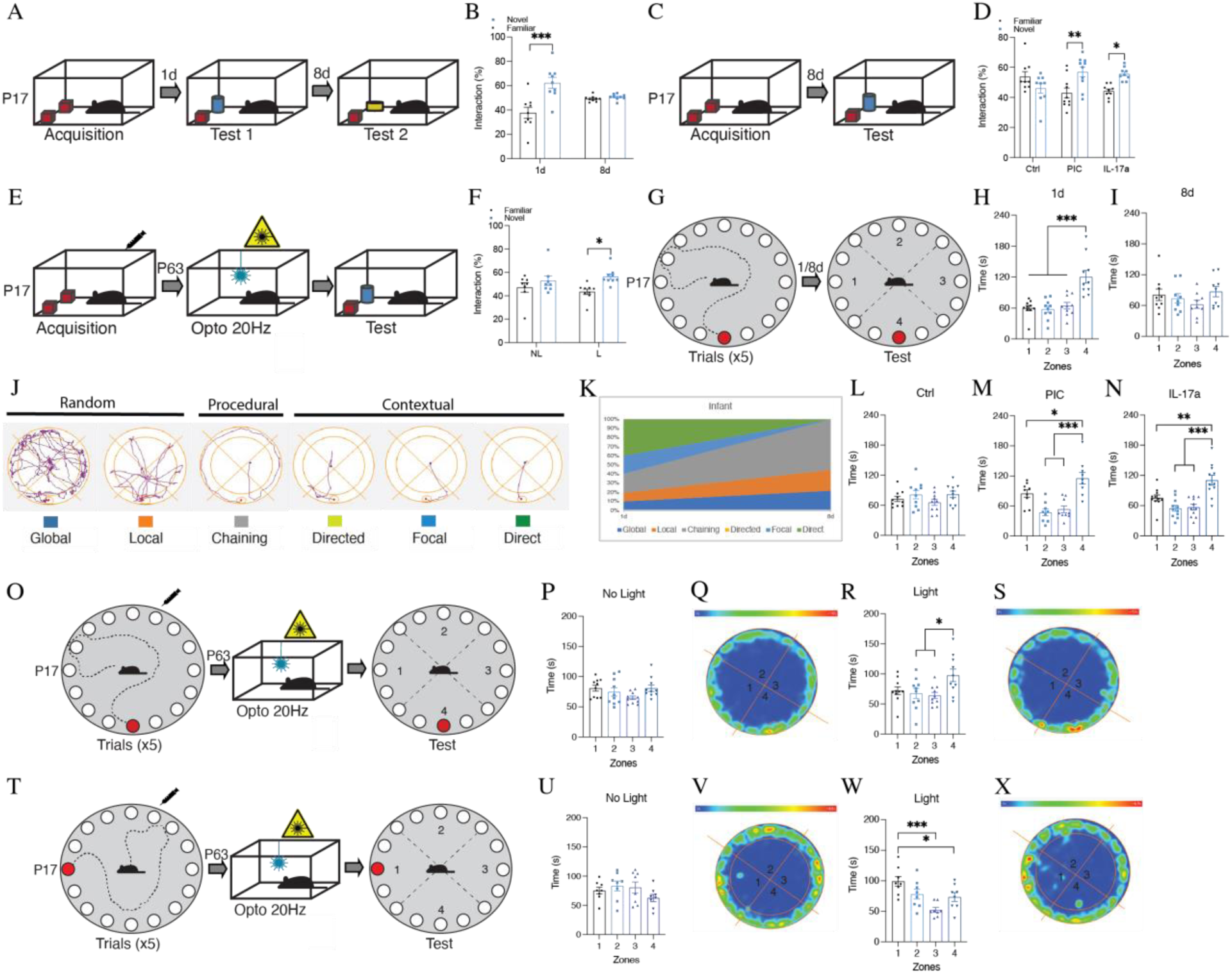
Infantile amnesia for context, object, and maze memory. (**A**, **C**) Behavioral schedule for novel object recognition. (**B**) Interaction index (%) of infant C57BLJ mice during novel object recognition test 1 or 8 d after acquisition (n=9). (**D**) Interaction index of MIA male infant offspring (n=9-10). (**E**) Behavioral schedule for DG optogenetic reactivation of an infant encoded engram for an object. (**F**) Interaction index of adult mice after optogenetic stimulation (n=9-10). (**G**) Behavioral schedule for the Barnes maze in infant C57BLJ mice. (**H**, **I**) Time spent in each zone during the test (**H**) 1 d or (**I**) 8 d after training. (**J**) Example representations of navigational search strategies. (**K**) Infant search strategies during testing 1 and 8 d after training. (**L**-**N**) Time spent in each zone by MIA offspring during test 8 d after training (n=9-11). (**O**, **T**) Behavioral schedule for DG optogenetic reactivation of an infant encoded engram for the Barnes maze. (**P**, **R**, **U**, **W**) Time spent in each zone during the test (n=8-9). *P < 0.05, **P < 0.01,*** P < 0.001 calculated by one-way ANOVA (**L**-**N**, **P**, **R**, **U**, **W**), two-way repeated measures ANOVA (**B**), or two-way ANOVA (**D**,** F**) with Bonferroni post hoc tests. Data presented as ±SEM.

We then investigated whether it was possible to rescue a ‘lost’ infant object memory by optogenetic stimulation of infant-labeled engram cells (Fig. 4E, SFig. 12F). On the test day, animals that received light stimulation 3 min before the test demonstrated a preference for the novel object (Fig. 4F). While animals that did not receive light stimulation before the test spent the same amount of time exploring both the novel and familiar object. These object memory engrams survive infantile amnesia, can be retrieved artificially, and their forgetting can be prevented by MIA.

Spatial and object tasks in rodents are useful for the development of parallel, translational object/context and maze learning assays in human behavior experiments. To expand our analysis of infantile amnesia to an active, maze-based task we probed memory for a spatial location in infant and adult C57BL/6J mice using a cued version of the Barnes maze (Fig. 4G). All cohorts sufficiently learned the escape hole’s location during training over five sessions, evidenced by a gradual decrease in the latency, and distance traveled, to reach the escape hole (SFig. 12G, H, K-M). Unlike adults, infant mice demonstrated memory retention for the location of the escape hole when tested 1, but not 8, days later (Fig. 4H, I, SFig. 12 I, J, N-S). To the best of our knowledge, this is the first demonstration of infantile amnesia using the Barnes maze paradigm. We assessed memory content profiling changes in navigational strategies taken by the animal during the task (Fig. 4J). Navigational strategies can be categorized as random, procedural, or contextual (*36–38*). Infant mice predominantly used contextual search strategies when tested for spatial memory on the Barnes maze 1 day after training (Fig. 4K). However, during the 8-day probe trial infant mice show a switch in navigational strategy, moving to procedural and random strategies (Fig. 4K). This adaptation in navigational search strategies reflects the forgetting of the location of the escape platform. In contrast, adult mice show similar search patterns both 1 and 8 days after training (SFig. 12T). Like other forms of memory, we also found that male offspring from MIA models do not show infantile amnesia for a spatial navigation task (Fig. 4L-N, SFig. 12U-W).

By labeling the engram ensembles for the location of the escape platform in the Barnes Maze in infants, we next investigated whether DG optogenetic stimulation of the engram in adulthood results in the location of the escape platform (Fig. 4O). The experimental light group received 3 min light stimulation before being placed on the Barnes maze for a probe test. The experimental light group spent significantly more time in the zone where the escape hole should be when compared to zone 2 and 3 (Fig. 4R). Although there was no significant difference in the total time spent between zone 4 and zone 1, heat map analysis of the mean time spent in each location showed a concentration around the location where the escape hole previously was located during training (Fig. 4S). No difference was seen in the total time spent between zones for the no-light control group (Fig. 4P, Q). Changing the location of the escape hole during memory encoding did not affect the recall of the location of the escape hole after light stimulation (Fig. 4T-X). Optogenetic stimulation of EYFP-expressing neurons in the DG (neurons labeled during infant encoding of the Barnes maze memory) allows for recall of the location of the escape hole in the Barnes maze.

To test whether the effect of immune activation on infantile amnesia is mediated by the downstream effector cytokine IL-17a, we next injected pregnant Il17a knockout (KO) dams with Poly I:C at E12.5 (Fig. 5A). Adult male and female offspring from MIA Il17a KO dams did not demonstrate repetitive behavior (SFig. 14E) or deficits in social behavior (SFig. 14A-D). Crucially, infant offspring show normal infantile amnesia for a CFC memory at P25 (Fig. 5B, C). Preventing maternal IL-17a signaling, using a genetic model, occluded the MIA effect on infantile amnesia in the offspring. Therefore, the behavioral effect seen following MIA is specifically dependent on IL-17a signaling. We investigated whether altering the timing of immune activation, from maternal to postnatal, would have the same effect on memory retention (Fig. 5D) (*39*). Postnatal immune activation in C57BLJ mice did not affect infantile amnesia, indicating that there is a critical window during prenatal development where IL-17a and immune activation can affect a mammal’s propensity for infantile amnesia (Fig. 5E-F). Induction of an inflammatory response in MIA offspring has been shown to temporarily rescue deficits in sociability in ASD (*25*). Therefore, we investigated whether we could reverse the effect of MIA on infantile amnesia in the offspring through the administration of IL-17a. MIA male offspring were injected with IL-17a three hours before the recall test at P25 (Fig. 5G). Regardless of whether injected with IL-17a or PBS, MIA male infant offspring showed normal memory retention for a CFC memory 8 days after training (Fig. 5H), indicating that the effect of MIA on infantile amnesia cannot be reversed through IL-17a administration.

**Fig. 5.**
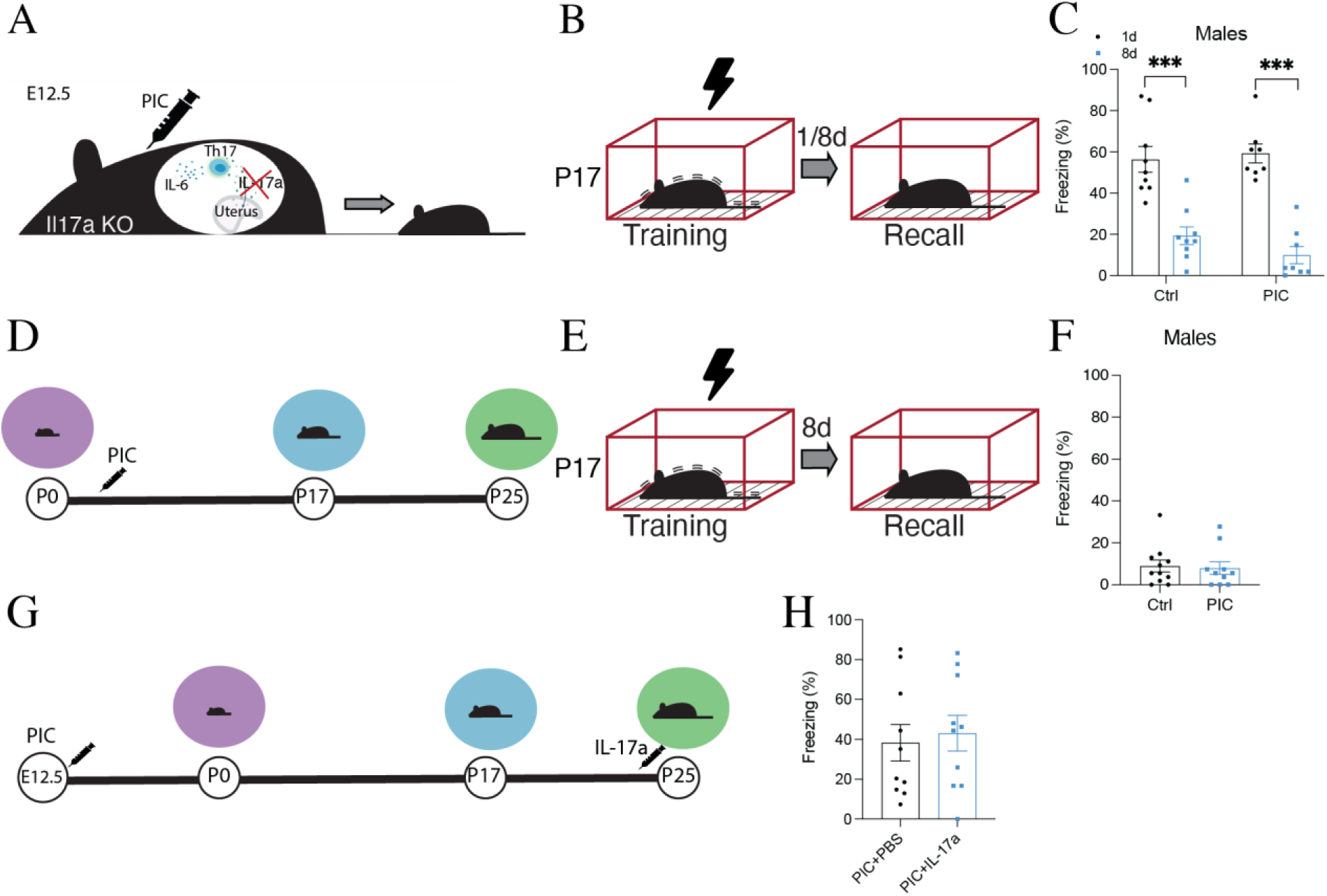
IL-17a is required for MIA effects on infantile amnesia. (**A**) Representative diagram of MIA in Il17a KO mice. (**B**) Behavioral schedule for CFC in Il17a KO male and female mice. (**C**, **D**) Freezing levels of male and femaleIl17a KO infant mice during recall 1 or 8 d after training. (**E**) Behavioral schedule for postnatal immune activation in C57BLJ infant mice. Syringe symbol represents Poly I: C injection at P3, P7, and P14. (**G**, **H**) Freezing levels of infant male and female mice when tested 8 d after training. (**I**, **J**) Behavioral schedule for IL-17a injection 3 h before recall at P25 in MIA C57bLJ offspring. (**K**) Freezing levels of infant male and female mice when tested 8 d after training. *P < 0.05, P < 0.01, P < 0.001* calculated by Student t-test (**F**, **H**) or two-way ANOVA (**C**) with Bonferroni post hoc tests. Data presented as ±SEM.

Together, these findings indicate that infantile amnesia for a wide range of memory types can be modulated by the immunological experiences of the subject during embryonic development. The effects of MIA preserve the natural retrievability of infant engrams across postnatal development. Our data shows that infant memories are suppressed during development, but that their engrams persist and can be acutely activated in adulthood by direct optogenetic stimulation (*23*). These findings demonstrate that optogenetic reactivation of an infant memory is not specific to CFC, memories for more complex navigational and recognition tasks can also be rescued. Furthermore, artificially updating an engram restored natural access to the target memory at both a behavioral and cellular level, reversing infantile amnesia. The specific connectivity patterns between engram cells, formed during infancy across distributed brain regions, also remain intact into adulthood and may account for stored information to be retained after infantile amnesia (*23*). Postnatal immune activation does not seem to affect infantile amnesia, making it plausibly driven by an innate predetermined switch. The brain state that permits infantile amnesia is absent in immunological models of ASD, and these findings are reminiscent of fly and human studies that have shown improved long-term retention of memories in cases of ASD (*40*, *41*). It is conceivable that the MIA offspring’s brain state may mirror that of precocial mammals, who do not show infantile amnesia (*4*, *42*). Infantile amnesia may represent a genetically canalized form of natural forgetting that can be modulated by developmental plasticity during critical periods. Future studies should determine the nature of the switching mechanism that determines when infantile amnesia occurs, its interaction with the immune system, and its reversible effect on engram cell function. Maternal immune activation models may offer new opportunities for translational memory studies across the lifespan in mice and humans (*43*).

## Acknowledgments

We thank R. Cusack and M. Ramaswami for useful discussions; J. O’Leary, M. Pezzoli, L. Marks, and other Ryan Lab members for collegial support and scientific input.

## Author contributions

Conceptualization: SDP, TJR

Methodology: SDP, COS, LL, TJR

Investigation: SDP, ES, LZ, COS, EPB

Funding acquisition: TJR

Project administration: SDP, TJR

Supervision: TJR

Writing – original draft: SDP, TJR

Writing – review & editing: SDP, ES, LZ, COS, LL, TJR

## Data and materials availability

All data are available in the main text or the supplementary materials.

## Supplementary Materials

### Materials and Methods

#### Subjects

The Ai32-FosTRAP mice were generated by crossing Fos-iCre (*27*, *28*) mice with Ai32(RCL-ChR2(H134R)/EYFP) (*29*, *30*) and selecting offspring carrying both the CreETR and EYFP transgenes. All mice were socially housed in numbers 2-5 littermates, on a 12 h light/dark cycle with access to food and water ad libitum. The day of birth was designated postnatal day 0 (P0). Infant (P17) mice were housed with parents until the time of weaning when they were group-housed by the same sex. Litters size was controlled, and no litter consisted of more than 6 pups. For infant experiments, each litter was counterbalanced across groups to limit litter effects. Each experimental group consisted of 4-5 liters. All mice were 7 weeks old before undergoing surgery. Post-operation mice were allowed to recover for a minimum of 2 weeks in their homecage before experimentation. All procedures relating to mouse care and treatment conformed with the health produces regulatory authority (HPRA) Ireland guidelines.

#### ChR2-EYFP expression in Ai32-FosTRAP mice

To label memory engram cells with this system, mice were injected intraperitoneally with 4-OHT (50mg/kg) 2 h post the learning event. The Fos-iCre line expresses the inducible c-Fos promoter. Injection of 4-OHT activates iCre recombinase. Activated iCre recombinase translocates to the nucleus and acts on two lox sites removing the stop codon that otherwise prevents expression of the ChR2/EYFP transgene. ChR2/EYFP transgene expression is driven by the pCAG promoter in iCre-expressing tissue 72 h after 4-OHT injection. For activity-dependent expression of ChR2-EYFP in Ai32-FosTRAP mice, Kainic acid (5mg/kg) was injected intraperitoneally followed by 4-OHT injection 2 h post kainic acid injection.

#### Stereotactic optical fiber implant

Mice were anesthetized using 500 mg kg-1 avertin. Each animal underwent bilateral craniotomies using a 0.5 mm diameter drill bit at −2.0 mm anteroposterior (AP), 1.3 mm mediolateral (ML) for DG injections. A bilateral patch cord optical fiber implant (200 core diameter) was lowered above the injection site (−1.9 mm DV). A layer of adhesive cement was applied to secure the optical fiber implant to the skull. Dental cement was applied to secure a protective cap to the implant and close the surgical site. Each animal received meloxicam analgesic (0.075 mL/5g) via subcutaneous injection and remained on a heating pad until fully recovered from anesthesia. All mice were allowed to recover for two weeks before any subsequent experiments. All fiber sites were verified histologically.

#### Immunohistochemistry

Mice were dispatched by overdose with 50 l sodium-pentobarbital and perfused transcardially with phosphate-buffered saline (PBS), followed by 4 % paraformaldehyde (PFA) in PBS. Extracted brains were kept in 4 % PFA at 4 oC overnight and stored in PBS. 50 □m coronal slices were cut using a vibratome and collected in PBS. Slices were washed in PBS-Triton X-100 (PBS-T) 0.2% followed by a 1 h blocking in PBS-T with 10% normal goat serum at room temperature before being incubated with the primary antibody at 4 oC overnight. Slices were washed in PBS-T 0.1 % followed by a 1 h incubation with the secondary antibody before undergoing another round of washing using PBS-T 0.1%. Vectashield DAPI was used to mount the slices onto super frost slides before visualization using an Olympus BX51 upright microscope. For cell counting experiments and dendritic spine analysis, an additional incubation with DAPI in PBS (1:1000) was carried out to label nuclei, and images were acquired using Leica SP8 gated stimulated emission depletion (STED) nanoscope. All images were taken at 40 X magnification.

#### Cell counting

To measure the extent to which populations of active cells overlap between the exposure to the same or different contexts, the number of EYFP+ and c-Fos+ immunoreactive neurons in the DG, AMG, RSC, and PAG were counted. All animals were sacrificed 45 min post-assay or optical stimulation for immunohistochemical analyses. Four coronal slices were taken from the dorsal hippocampus of each animal. For optogenetic stimulation experiments, only slices with accurate implant location sites were used for counting. Sections were imaged on a Leica SP8 gated stimulated emission depletion (STED) nanoscope at a magnification of 40X. The area of interest was manually identified and the area of each region of interest was calculated using Fiji. To calculate the total number of DAPI cells in the DG, the average diameter of a sample of EYFP-positive cells in the DG was taken from each animal and the area of the cell was calculated. The total number of cells was estimated as the area of the DG divided by the area of the cell. For the AMG, PAG, and RSC, the number of DAPI cells in 3 randomly selected regions of interest were counted and used along with the total area of the region to determine the total number of DAPI cells. The total number of c-Fos positive and EYFP positive cells were manually identified and counted for each region using Adobe Photoshop CC 2018. To calculate the percentage of cells expressing EYFP in each region, the total number of EYFP-positive cells was divided by the total number of DAPI-positive cells for each area. To calculate the percentage of cells expressing c-Fos in each region, the total number of c-Fos positive cells was divided by the total number of DAPI-positive cells for each area. To quantify engram reactivation, the number of EYFP and c-Fos positive cells were quantified as a percentage of total EYFP or DAPI positive cells.

#### Dendritic spine analysis

The dendrites of DG engram cells in Ai32-FosTRAP mice were analyzed after recall at P25. Dendritic spines were imaged using the Leica SP8 gated STED scope. Z-stack images (10 μm) of 50 μm coronal brain sections were taken using Leica Application Suite X (LasX) software (line average 2, zoom factor 1.7-1.9) at 40X. Imaris software (Oxford Instruments, Imaris v9.5) was used to carry out dendritic spine analysis. Dendrites were traced (10 μm) using a semi-automated neurofilament tracer tool and dendritic spines were individually highlighted and manually traced with the software. Each fragment represents a different ChR2-EYFP+ cell. Parameters were generated automatically using Imaris software.

#### Behavioral assays

All behavioral experiments were conducted during the light cycle of the day (07:00 am to 7:00 pm). All behavioral subjects were individually habituated to handling by the investigator for 3 min on 3 consecutive days. Handling took place in the breeding room. Immediately before each handling session mice were transported in separate cages to and from the vicinity of the experimental room to habituate them to the journey.

#### Contextual fear conditioning

Three distinct contexts were used for contextual fear conditioning (CFC). Context A was a 31 x 24 x 21 cm Med Associates chamber with a removable grid floor (bars 3.2 mm diameter, spaced 7.9 mm apart), opaque triangular ceiling, and scented with 0.25 % benzaldehyde. Context B chambers were 29 x 25 x 22 cm Coulbourne Instruments chambers with Perspex white floors, bright white lighting, and scented with 1 % acetic acid. All conditioning sessions were conducted in Context A. Each session was 330 s in duration with three 0.75 mA shocks of 2 s duration delivered at 150 s, 210 s, and 270 s. Mice were removed from the conditioning chamber 60 s following the final footshock and returned to their homecage. All recall or context generalization tests were 180 s in duration. Testing conditions were identical to training conditioning except that no shocks were presented. Up to 4 mice were run simultaneously in the 4 identical chambers. Floors of chambers were cleaned with Trigene before and between runs. Mice were transported to and from the experimental room in separate Perspex cages. All experimental groups were counter-balanced for chambers within contexts.

#### Contextual pre-exposure

For context pre-exposure mice were exposed to Context A (31 x 24 x 21 cm Med Associates chamber with a removable grid floor) for 10 min. Mice were removed from the chamber and placed back in their homecage. The following day, mice were placed back in Context A where an immediate foot shock (1mA, 2s) was delivered. Mice were removed from the chamber 1 min after the shock and placed back in their homecage. Up to 4 mice were run simultaneously in the 4 identical chambers except for during the update when one mouse was run at a time. All recall or context specificity tests were 180 s in duration. Floors of chambers were cleaned with Trigene before and between runs.

#### Novel object recognition

Each subject was allowed to habituate to the rectangular testing arena for 10 min on the day before object acquisition. During object acquisition, two identical objects were positioned on adjacent walls of the arena. Mice were allowed to explore both objects freely for a 10 min period before being placed back in their homecage. On testing, one familiar object was replaced with a novel object and each subject was introduced for a 5 min exploration period. Object exploration was defined as the time when the subject’s nose came within a 2 cm radius of the object. In between each trial, the testing arena and objects were cleaned with Trigene. For optogenetic reactivation experiments, animals received 3 min of blue light stimulation directly before being placed into the testing arena.

#### Barnes maze

Training consisted of five trials with a 10 min limit per trial. At the start of each trial, the mouse was placed in a black PVC start chamber located in the center of the maze apparatus. After 15 secs, the start chamber was removed, and the latency and distance traveled to enter the escape tunnel were recorded. Subjects were placed back in their homecage and allowed 40 min in between each trial. The position of the escape tunnel remained in a fixed location relative to spatial cues in the room. For each probe trial, the escape tunnel was removed from the apparatus. Each mouse was allowed 5 min to freely explore the four quadrants of the maze before being removed and placed back in its homecage. The latency and distance traveled before reaching the location of the removed escape tunnel, along with the time spent in each quadrant were recorded. The apparatus and escape tunnel were cleaned with Trigene after each trial. For optogenetic reactivation experiments, animals received 3 min of blue light stimulation directly before being placed into the testing arena.

#### Three-chamber social interaction

Mice were habituated to the chamber with two empty holders for 10 min. The next day, mice were placed in the middle of the chamber and allowed to explore the three chambers for a period of 5 min. During this testing period, a social object (novel mouse) was contained in one holder in one chamber and an inanimate object (lego blocks) was contained in a holder in the other chamber. Object exploration and distance traveled were tracked using ANY-maze video tracking software.

#### Repetition assay

A large homecage was filled 5 cm deep with wood chipping bedding and lightly tamped down to make an even flat surface. A consistent pattern of 20 identical glass marbles (15mm diameter) was evenly placed (4 cm apart) on the surface of the wood chip bedding. Mice were left alone in the testing arena for 30 min. A picture was taken before and after the test for analysis. A marble was considered buried if 2/3rds of the depth of the marble was buried.

#### Optogenetic engram reactivation

Optogenetic engram reactivation was carried out in Context C, a 31 x 24 x 21 cm Med Associates chamber with a white Perspex floor and curved insert, a dim light level and scented with 0.25 % acetophenone. The optical fiber implant was connected to a 45 nm laser diode fiber light source (Doric LDFLS 450/080). Habituation sessions lasted for 12 min. During testing, the 12 min session was divided into four 3 min epochs split into two light-off and two light-on epochs. During the light-on epochs, blue light stimulation (20 Hz) with a pulse width of 15 ms was delivered through the optical fiber implant for the entire 3 min duration. At the end of the 12 min session, the mouse was immediately detached from the laser and returned to its homecage.

#### Artificial updating

Context A was a 31 x 24 x 21 cm Med Associates chamber with a removable Perspex black floor, black triangular ceiling, and scented with 0.25 % benzaldehyde. Context B exposure was carried out in a rectangular testing arena. Artificial updating of a contextual memory was carried out in Context C, a 29 x 25 x 22 cm Coulbourne Instruments chamber with removable grid floors, bright white lighting, and scented with 1 % acetic acid. Contexts A, B, and C were all located in different behavioral rooms. The optical fiber implant was connected to a 45 nm laser diode fiber light source (Doric LDFLS 450/080). The 7 min session consisted of a 2 min acclimatization period followed by 5 min of blue light stimulation (20Hz) with a pulse width of 15 ms delivered through the optical fiber implant for the entire 5 min duration. Three shocks, 0.75 mA of 2 s duration, were delivered at minutes 4, 5, and 6 of the session. At the end of the 7 min session, the mouse was immediately detached from the laser and returned to its homecage.

#### Maternal immune activation

Mice were mated overnight for one night. Females were weighed and checked for seminal plugs, noted as E0.5. Pregnant dams were injected SC with a single dose of Poly(I: C) (Sigma Aldrich) at 20mg/kg, rmlIL-17A (Immunotools) at 50ug/kg, or control PBS on E12.5. For rmlIL-17A injections at P25, mice were injected SC with a single dose of rmlIL-17A (Immunotools) at 50ug/kg or control PBS 3 h before the recall test.

#### Postnatal immune activation

Mice were injected SC with a single dose of Poly(I: C) (Sigma Aldrich) at 20mg/kg or control PBS on P3, P7, and P14.

#### Genotyping

Genomic DNA was extracted from the ear punches of each mouse and sent to Transnetyx for genotyping.

#### Analysis and statistics

All experiments were analyzed blind to the experimental group. All videos were randomized before manual scoring. Behavioral performance was recorded by a digital video camera. Videos were manually scored individually, and investigators were blind to the experimental condition and test day during all manual scoring. Data analysis and statistics were conducted using GraphPad Prism 6.00 (GraphPad software). Unpaired Student’s t-tests were used for independent group comparisons. Paired Student t-tests were used to assess differences within groups. ANOVA followed by a Bonferroni post hoc test was used to determine conditions that were significant from each other where appropriate. Outliers were detected using Grubbs’ test where p < 0.05. For cell counting analysis results were analyzed per mouse using nested t-test or nested ANOVA followed by Bonferroni post hoc test. Results are displayed as mean with SEM and deemed significant when p < 0.05.

---

**Fig. S1.**
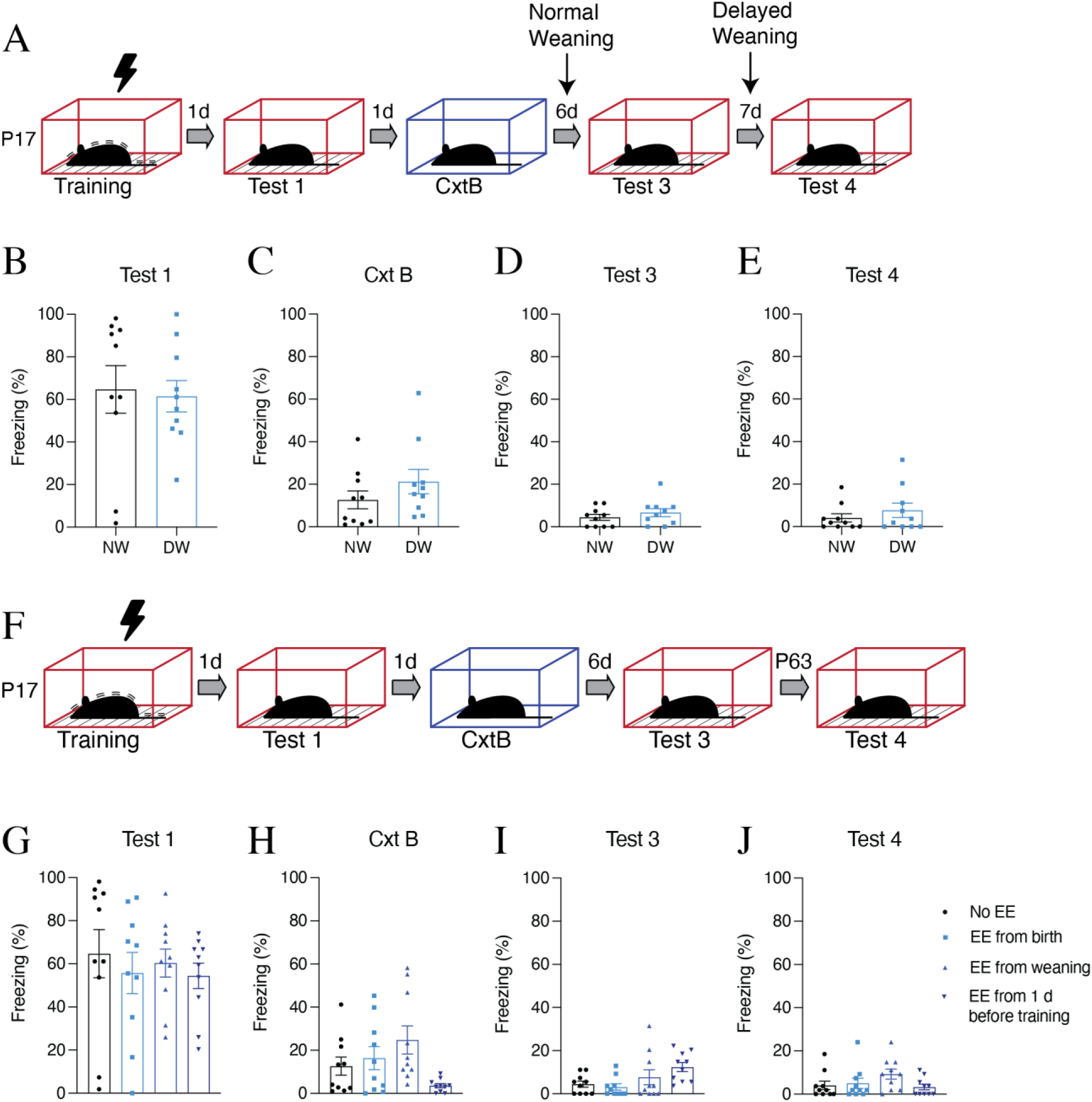
Delayed weaning or environmental enrichment does not affect infantile amnesia. (**A**) Behavioral schedule for CFC in infant (P17) C57BLJ male mice subjected to normal (P21) or delayed (P24) weaning. The lightning symbol represents footshocks. (**B**-**E**) Freezing levels during recall tests (n=10) at Test 1 (P18), CxtB (P19), Test 3 (P25), and Test 4 (P32). (**F**) Behavioral schedule for CFC in infant C57BLJ male mice. (**G**-**J**) Freezing levels during recall tests (n=10). *P < 0.05, P < 0.01, P < 0.001* calculated by Student t-test (**B**-**E)** or one-way ANOVA (**G**-**J**) with Bonferroni post hoc tests. Data presented as ±SEM.

**Fig. S2.**
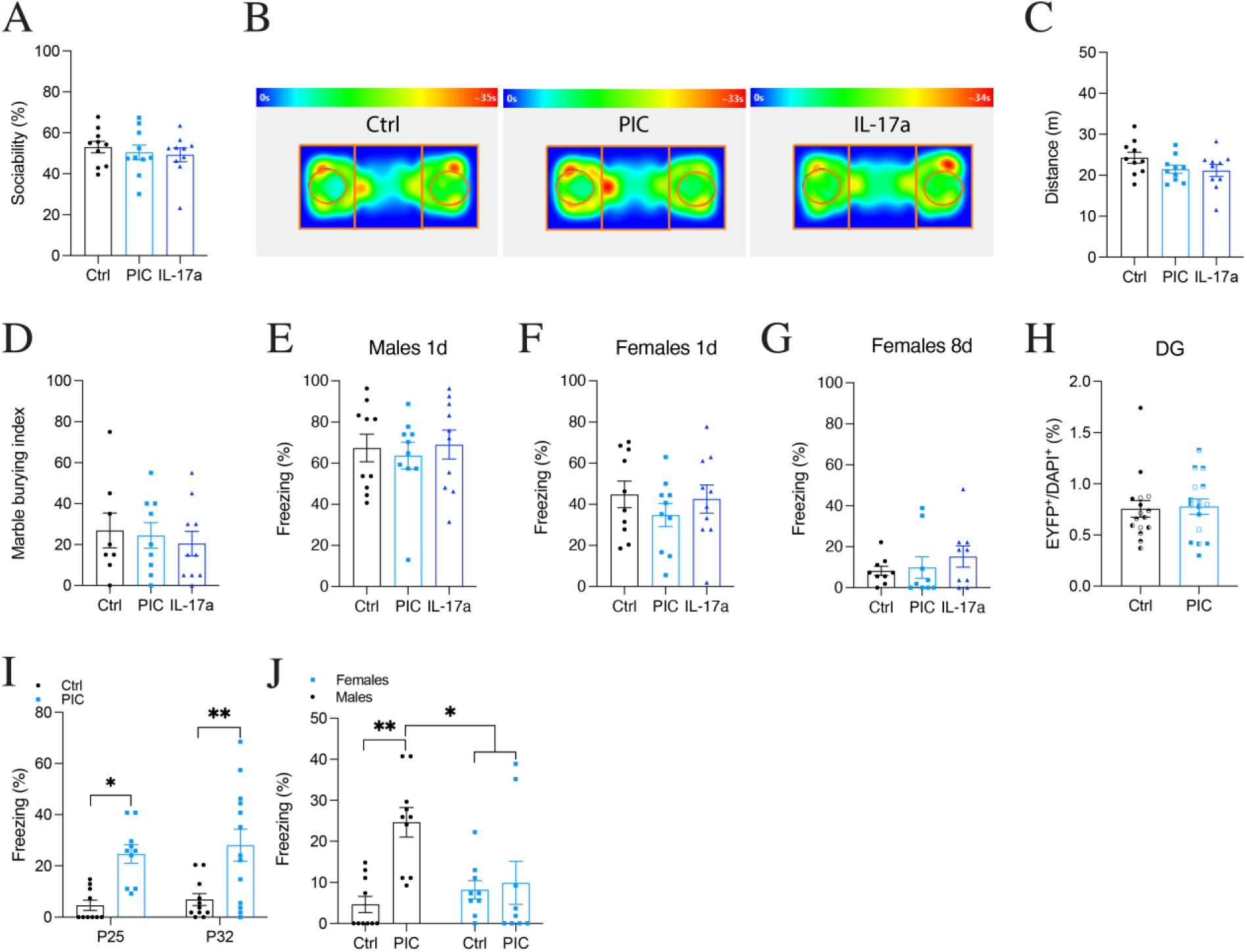
MIA female offspring do not demonstrate social interaction deficits or repetitive behavior but do show infantile amnesia for a fear memory. (**A**) Social interaction index (%) of adult (P63) C57BLJ female MIA offspring (n=10). (**B**) Heat map analysis of time spent in each chamber during social interaction task. (**C**) Total distance (m) traveled during social interaction test (n=10). (**D**) Number of marbles buried by adult female MIA offspring during marble burying task (n= 8-10). Memory recall for CFC in context A for male MIA offspring testing 1 d after training (n=10). (**F**, **G**) Recall for CFC in context A for female MIA offspring tested (**F**) 1 d (n=9-10) or (**G**) 8 d (n= 9) after training. (**H**) ChR2-EYFP^+^ cell counts in the DG (N= 4, n=4) of female MIA offspring after recall at P25. (**I**) Memory recall for CFC in context A for male MIA offspring tested 8 or 15 d after training n=11 (Ctrl), n=13 (PIC). (**J**) Memory recall comparison in male and female MIA offspring 8 d after training. *P < 0.05, P < 0.01, P < 0.001* calculated by nested Student t-test (**H**), one-way ANOVA (**A**, **C**-**F**), or two-way ANOVA (**I**) with Bonferroni post hoc tests. Data presented as ±SEM.

**Fig. S3.**
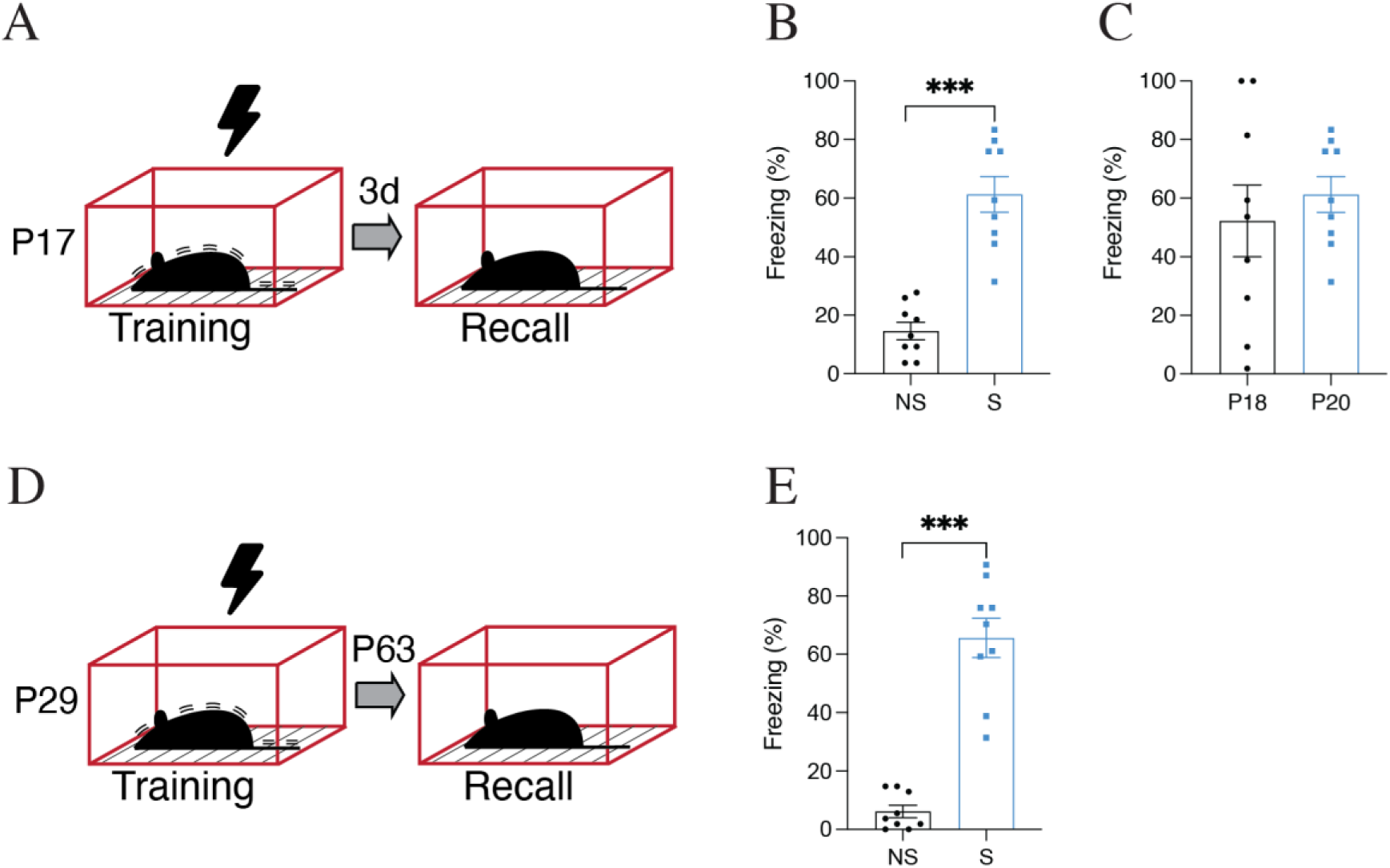
Infant mice show similar memory retention when tested 1 or 3 d after training at P17. (**A**) Behavioral schedule for CFC in C57BLJ P17 mice. The syringe symbol represents 4-OHT injection 2 h after exposure. The black lightning symbol represents foot shocks. (**B**, **C**) Freezing levels (%) during recall in context A. (**D**) Behavioural schedule for CFC in C57BLJ P29 mice. (**E**) Freezing levels (%) during recall in context A at P63. *P < 0.05, P < 0.01, P < 0.001* calculated by Student t-test (**B**, **C**, **E)**. Data presented as ±SEM.

**Fig. S4.**
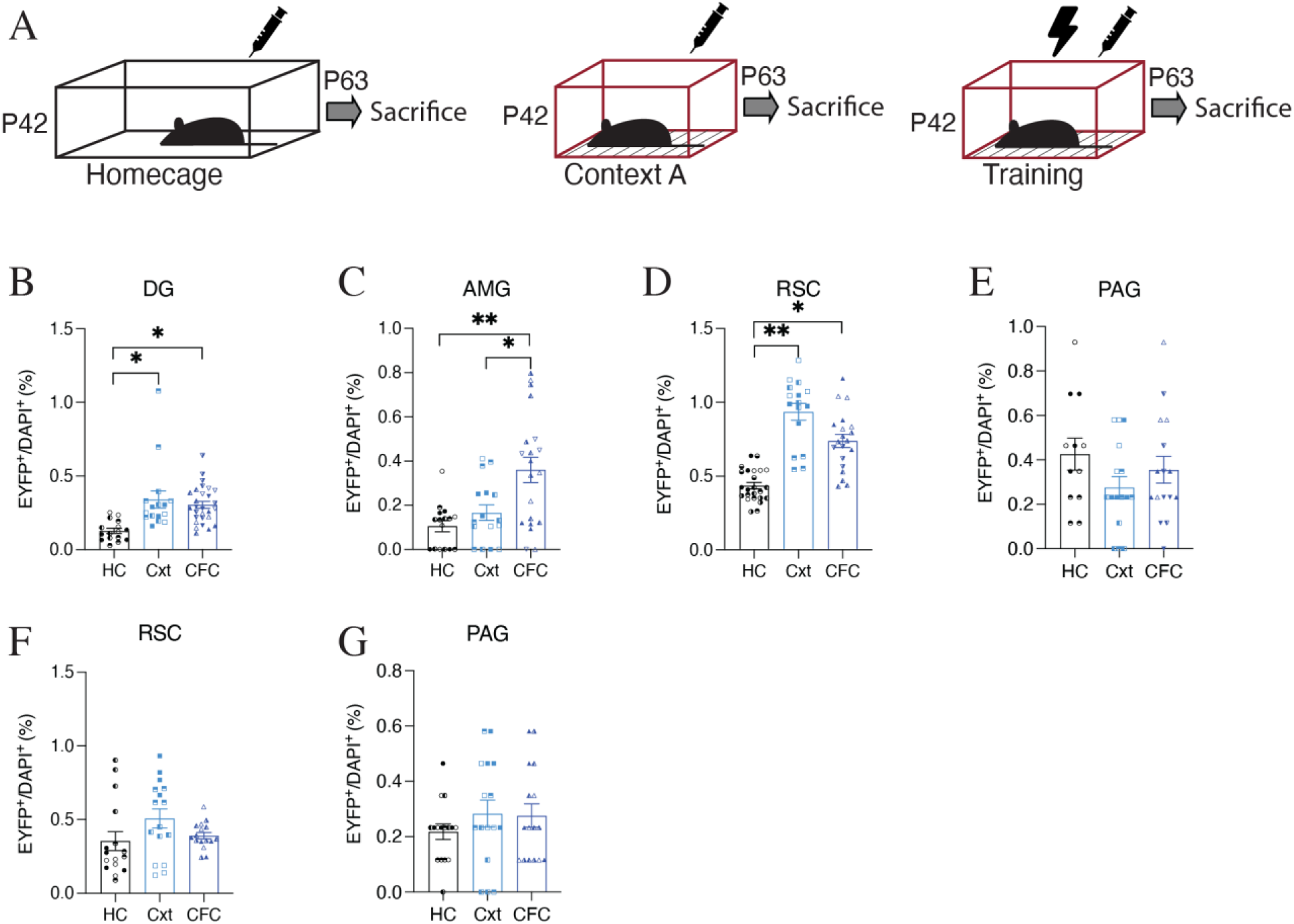
Adult Ai32-FosTRAP mice show activity-dependent increases in ChR2-EYFP labeling. (**A**) Behavioral schedule. The syringe symbol represents 4-OHT injection 2 h after exposure. The black lightning symbol represents foot shocks. (**B**-**E**) Quantification of ChR2-EYFP^+^ cells in the (**B**) DG (N=4-7, n=4), (**C**) AMG (N=4-5, n=4), (**D**) RSC (N=4-6, n=4) and (**E**) PAG (N=3-4, n=4) for an adult (P42) encoded homecage (HC), context (Cxt) and contextual fear (CFC) memory. Exposure to a context resulted in significantly more ChR2-EYFP^+^ cells in the (**B**) DG (p < 0.001) and (**D**) RSC (p < 0.001). Quantification of ChR2-EYFP^+^ cells after CFC was significantly higher in the (**B**) DG (p < 0.05), (**C**) AMG (p < 0.001), and the (**D**) RSC (p < 0.001). (**F**, **G**) ChR2-EYFP^+^ cell counts (N=4, n=4) in the RSC and PAG for an infant (P17) encoded HC, Cxt, and CFC memory. *P < 0.05, P < 0.01, P < 0.001* calculated by nested one-way ANOVA (**B**-**G**) with Bonferroni post hoc tests. Data presented as ±SEM.

**Fig. S5.**
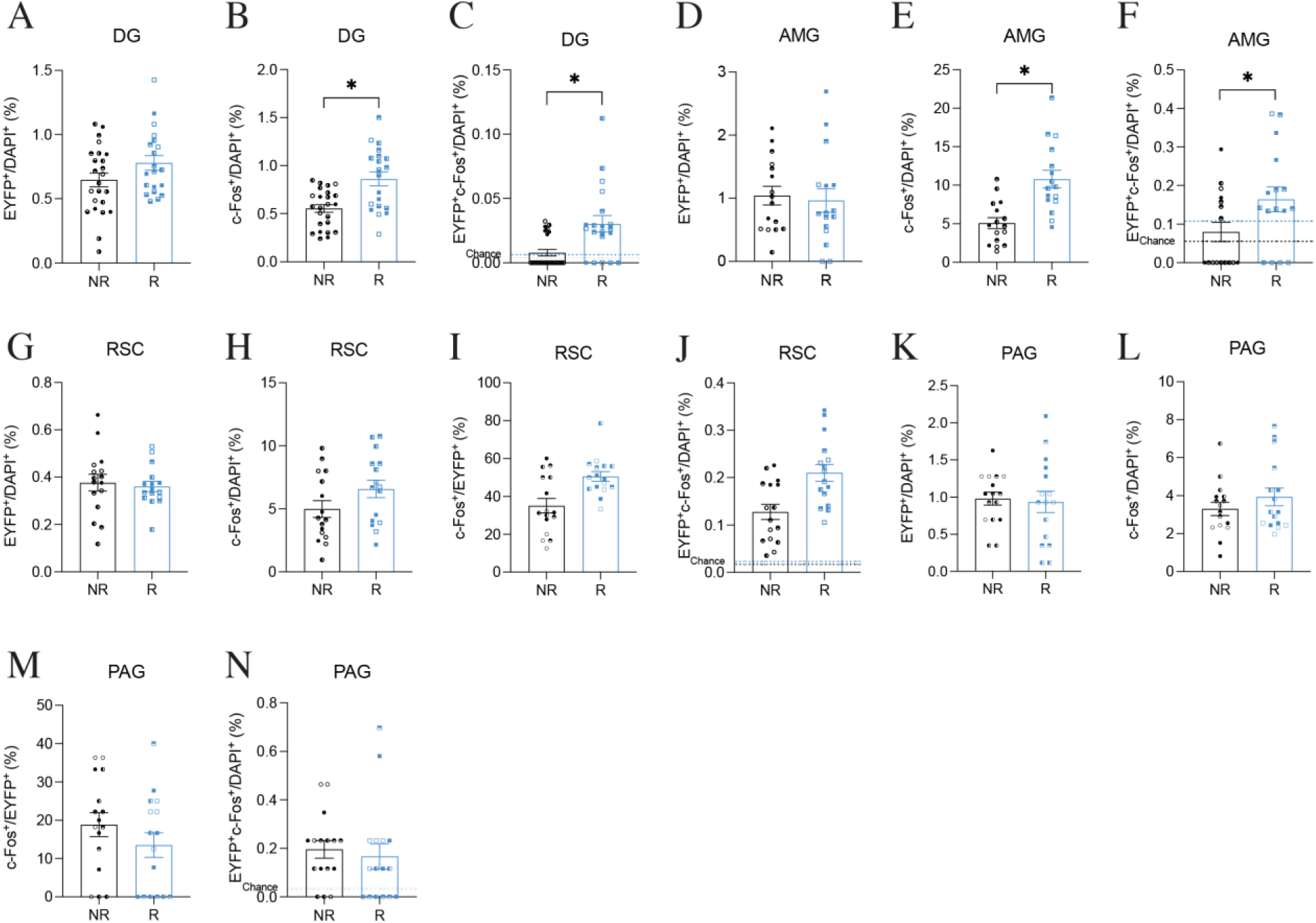
Engram reactivation is increased in the DG and AMG after natural recall at P20. (**A**-**N**) Histological analysis of cell counts after recall at P20. Percentage of (**A**) ChR2-EYFP^+^ and (**B**) c-Fos^+^ cells in the DG (N=5-6, n=4). There was a significant increase in c-Fos^+^ cells in the DG after recall at P20 (p < 0.05). (**C**) Percentage of DAPI cells positive for both ChR2-EYFP and c-Fos. (**C**) Engram reactivation as a percentage of DAPI^+^ was significantly higher in the DG after recall at P20 (p < 0.05). Percentage of (**D**) ChR2-EYFP^+^ and (**E**) c-Fos^+^ cells in the AMG (N=4, n=4). There was a significant increase in c-Fos^+^ cells in the AMG after recall at P20 (p < 0.05). (**F**) Percentage of DAPI cells positive for both ChR2-EYFP and c-Fos. (**F**) Engram reactivation as a percentage of DAPI^+^ in the AMG was significantly higher after recall at P20 (p < 0.05). Percentage of (**G**) ChR2-EYFP^+^ and (**H**) c-Fos^+^ cells in the RSC (N=4, n=4). Engram reactivation in the RSC as a percentage of (**I**) ChR2-EYFP^+^ and (**J**) DAPI^+^. Percentage of (**K**) ChR2-EYFP^+^ and (**L**) c-Fos^+^ cells in the PAG (N=4, n=4). Engram reactivation in the PAG as a percentage of ChR2-EYFP^+^ (**M**) and (**N**) DAPI^+^. *P < 0.05, P < 0.01, P < 0.001* calculated by nested t-test (**A**-**N**). Data presented as ±SEM.

**Fig. S6.**
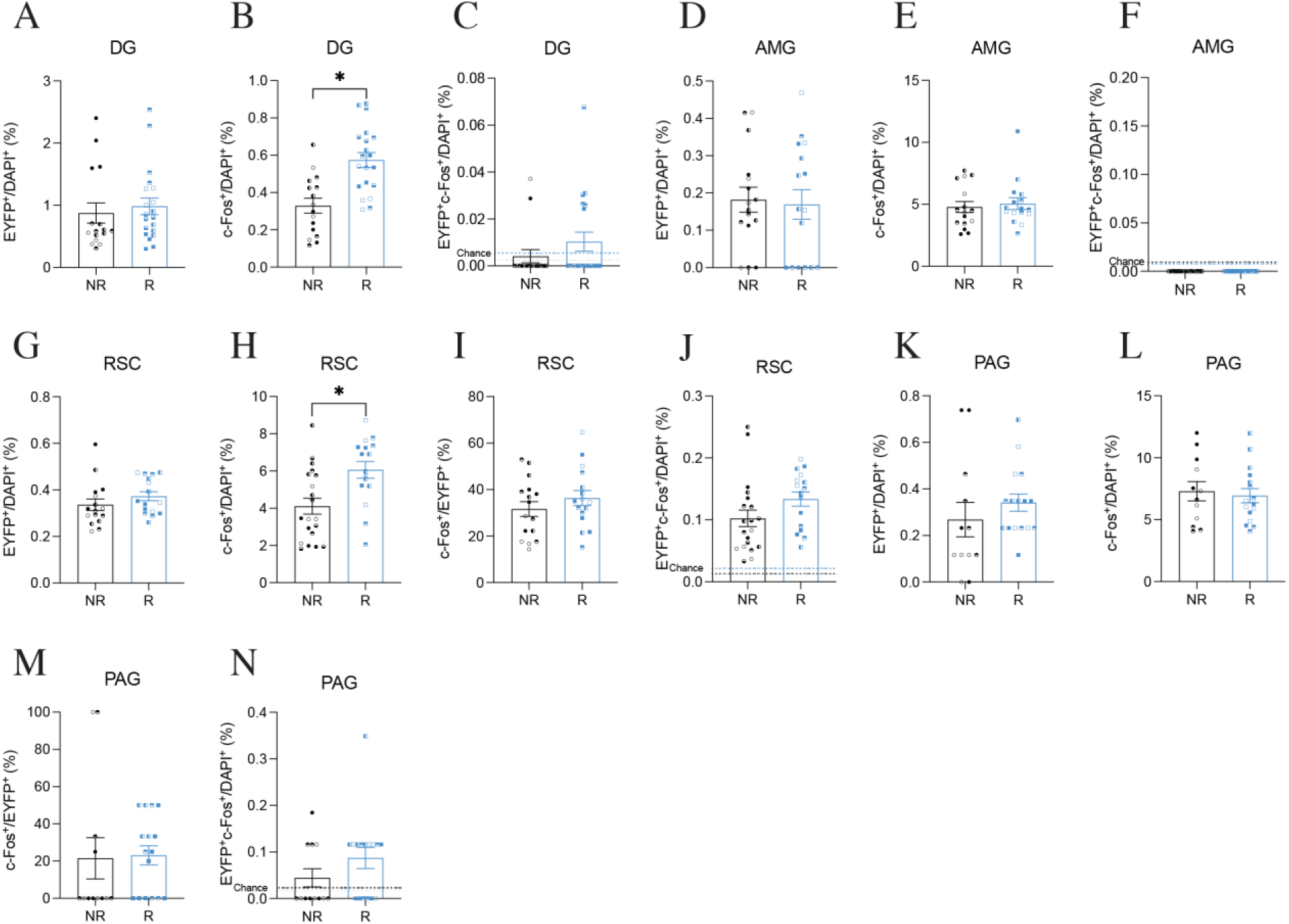
Presentation of recall cues for an infant encoded context does not result in engram reactivation at P63. (**A**-**N**) Histological analysis of cell counts after recall at P63. Percentage of (**A**, **D**) ChR2-EYFP^+^ and (**B**, **E**) c-Fos^+^ cells in the DG (N=4-5, n=4) and AMG (N=4, n=4). (**B**) There was a significant increase in c-Fos^+^ cells in the DG after recall at P63 (p < 0.05). (**C**, **F**) Engram reactivation as a percentage of DAPI^+^ in the DG and AMG. Percentage of (**G**, **K**) ChR2-EYFP^+^ and (**H**, **L**) c-Fos^+^ cells in the RSC (N=4-5, n=4) and PAG (N=3-4, n=4). (**H**) There was a significant increase in c-Fos^+^ cells in the RSC after recall at P63 (p < 0.05). Engram reactivation in the RSC and PAG as a percentage of (**I**, **M**) ChR2-EYFP+ and (**J**, **N**) DAPI^+^. *P < 0.05, P < 0.01, P < 0.001* calculated by nested t-test (**A**-**N**). Data presented as ±SEM.

**Fig. S7.**
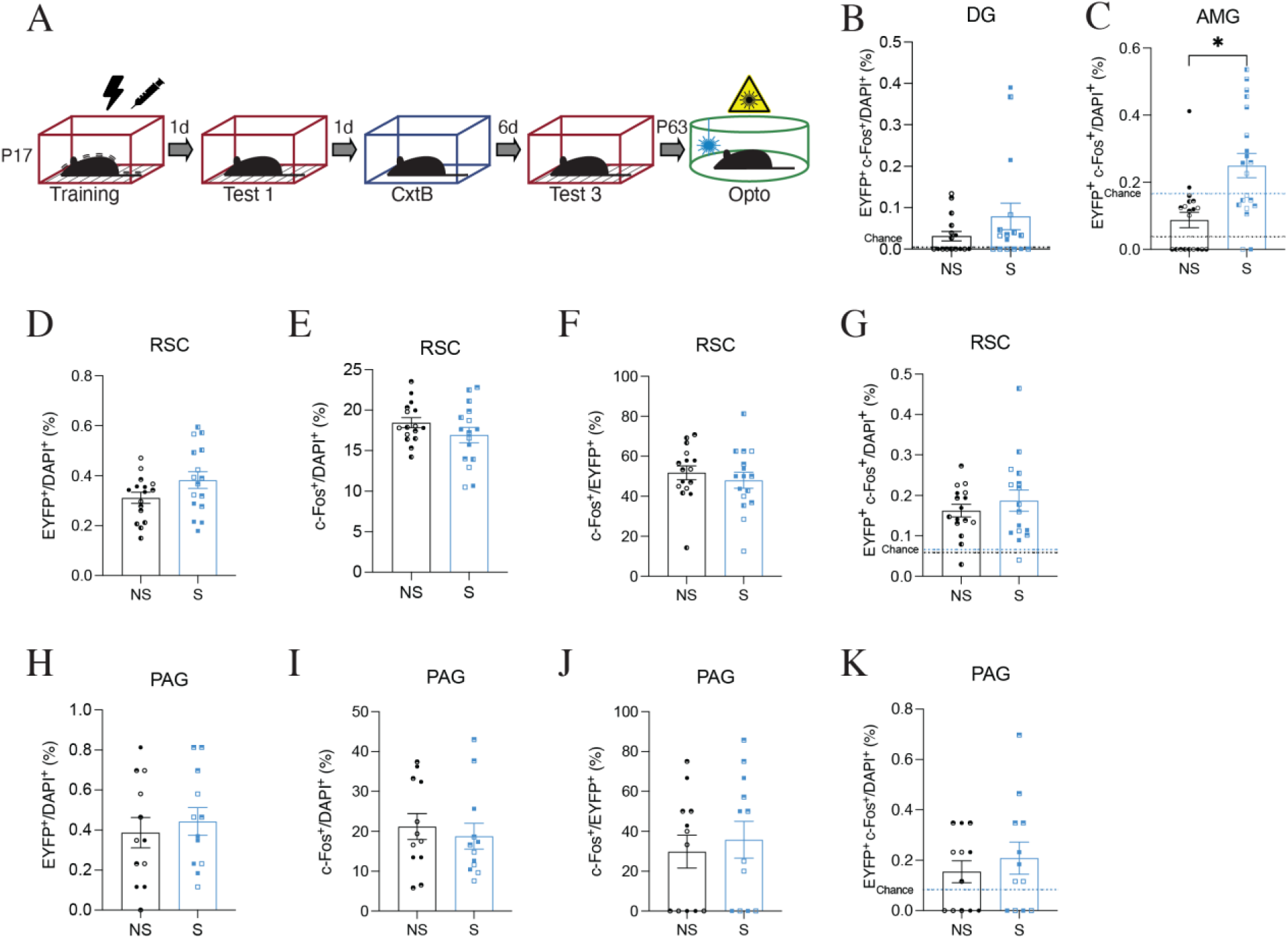
Optogenetic reactivation of an infant encoded engram. (**A**) Behavioral paradigm. (**B**-**K**) Cell counts after optogenetic reactivation of an infant encoded engram in adult Ai32-FosTRAP mice (Fig. 3A). (**D**, **E**) Percentage of DAPI cells positive for both ChR2-EYFP and c-Fos in the DG and AMG. (**C**) Engram reactivation as a percentage of DAPI^+^ cells was significantly higher (p < 0.05) in the AMG (N=5, n=4) after optogenetic stimulation of an infant-encoded fear memory. Percentage of (**D**) ChR2-EYFP^+^ and (**E**) c-Fos^+^ cells in the RSC (N=4, n=4). Engram reactivation in the RSC as a percentage of (**F**) ChR2-EYFP^+^ and (**G**) DAPI^+^. Percentage of (**J**) ChR2-EYFP^+^ and (**K**) c-Fos^+^ cells in the PAG (N=4, n=3-4). Engram reactivation in the PAG as a percentage of (**K**) ChR2-EYFP^+^ and (**K**) DAPI^+^. *P < 0.05, P < 0.01, P < 0.001* calculated by nested t-test (**B**-**K**). Data presented as ±SEM.

**Fig. S8.**
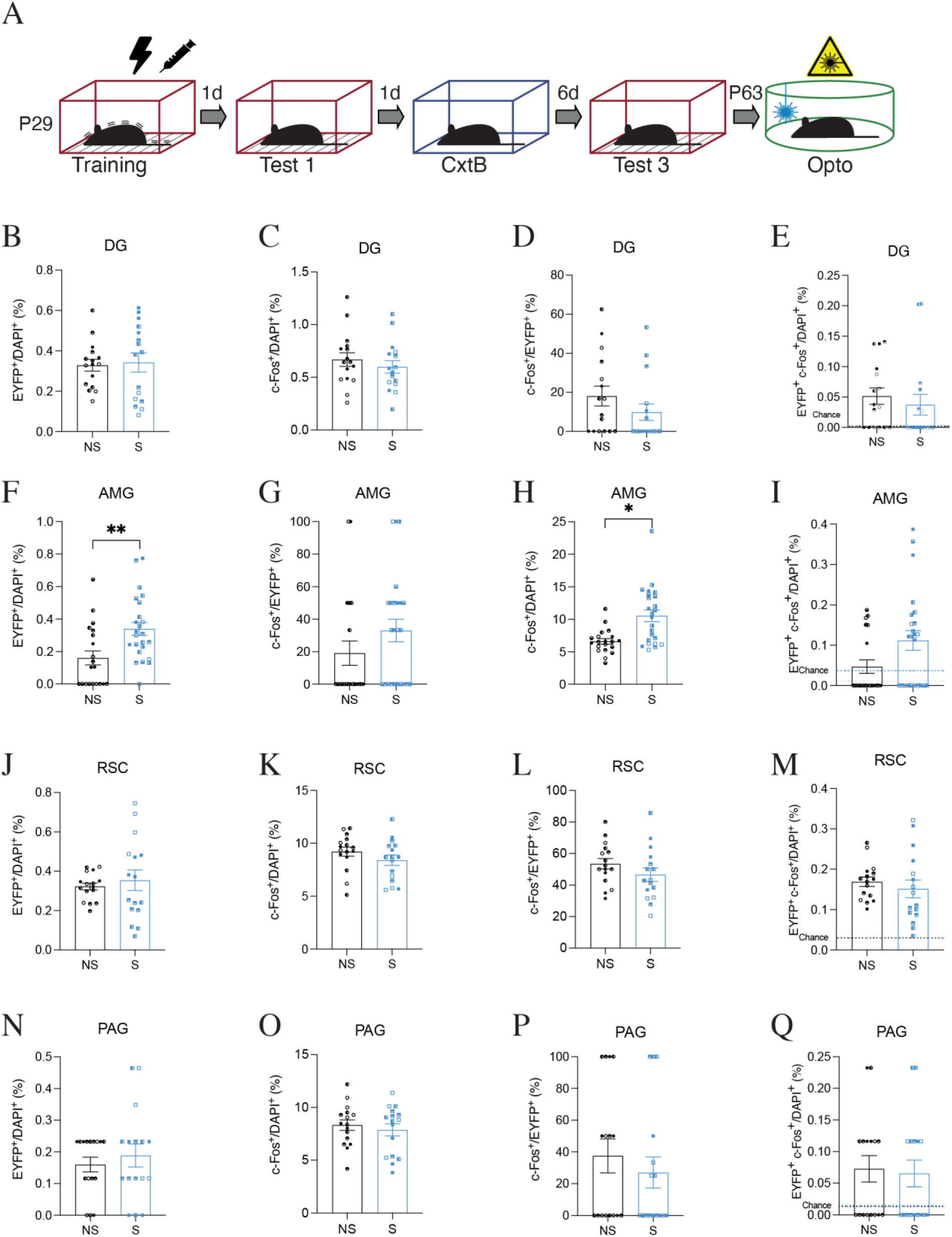
Optogenetic stimulation of an CFC engram encoded after the infantile period causes an increase in activity in the amygdala in adult mice. (**A**) Behavioral schedule for optogenetic reactivation of DG engram cells, labeled at P29, in adult Ai32-FosTRAP mice. The black lightning symbol represents foot shocks. The syringe symbol represents 4-OHT injection 2 h after training. Percentage of ChR2-EYFP^+^ cells labeled during training in the (**B**) DG (N=4, n=4), (**F**) AMG (N=5-6, n=4) (p < 0.05), (**J**) RSC (N=4, n=4), and (**N**) PAG (N=4, n=4). Percentage of c-Fos^+^ cells in the (**C**) DG, (**G**) AMG, (**K**) RSC, and (**O**) PAG after optogenetic stimulation at P63. Engram reactivation as a percentage of (**D**, **H**, **L**, **P**) EYFP^+^ and (**E**, **I**, **M**, **Q**) DAPI^+^ cells. *P < 0.05, P < 0.01, P < 0.001* calculated by nested t-test (**B**-**Q**). Data presented as ±SEM.

**Fig. S9.**
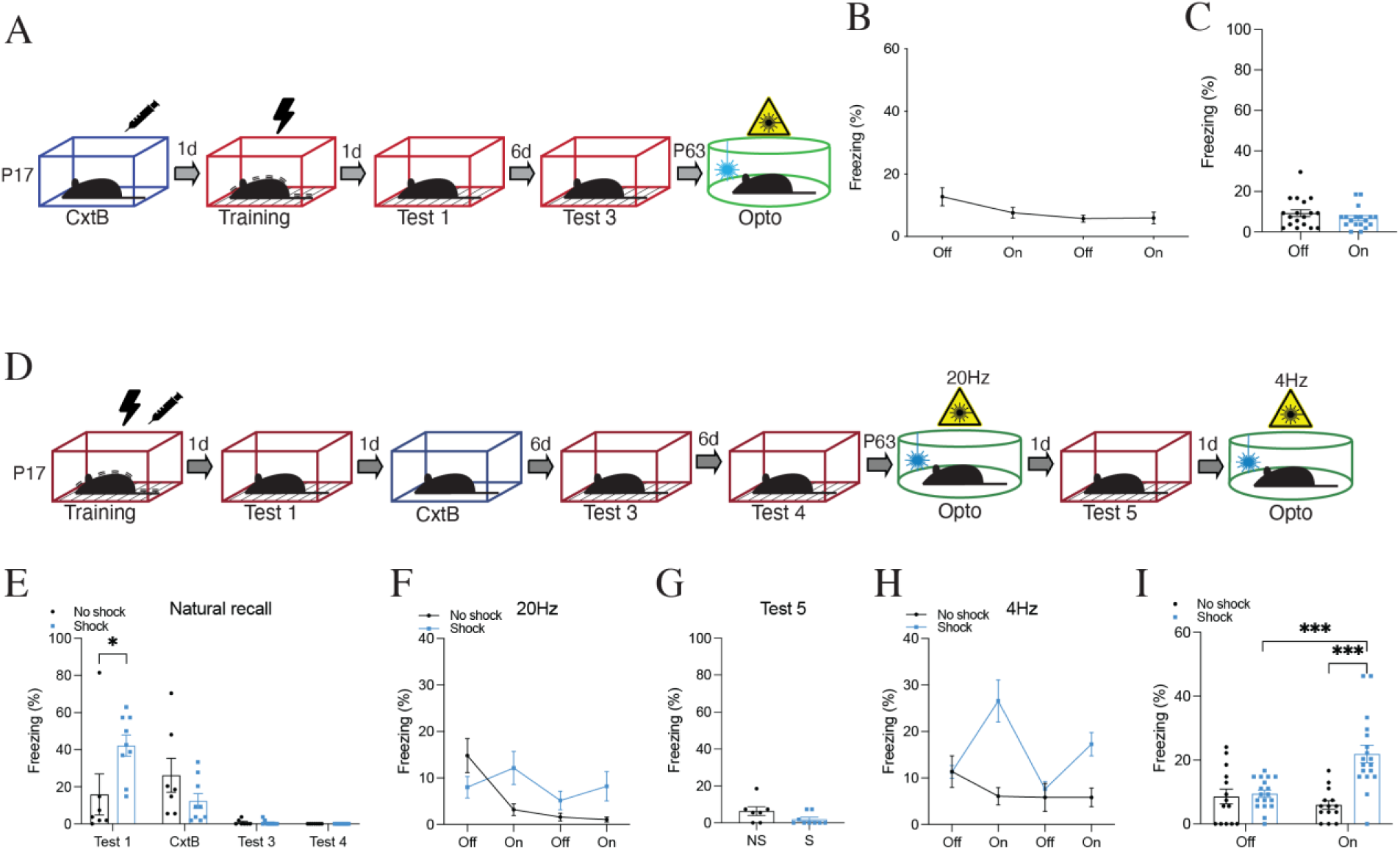
Optogenetic reactivation of an infant engram in the CA1 elicits freezing behavior in adult mice. (**A**) Behavioral schedule for engram reactivation of a neutral context B in the DG of Ai32-FosTRAP male mice. The black lightning symbol represents foot shocks. The syringe symbol represents 4-OHT injection 2 h after context exposure. (**B**) Memory recall in context C (engram reactivation) with light-off and light-on epochs (n=9). (**C**) Freezing for the two light-off and light-on epochs averaged. (**D**) Behavioural schedule for infant engram reactivation in the CA1 of adult Ai32-FosTRAP male mice n=7 (NS), n=9 (S). (**E**, **G**) Freezing levels during natural memory recall. (**F**, **H)** Memory recall in context C (engram reactivation 20 Hz (**F**) or 4 Hz (**H**)) with light-off and light-on epochs. (**I**) Freezing for the two light-off and light-on epochs during 4 Hz stimulation averaged. *P < 0.05, P < 0.01, P < 0.001* calculated by Student t-test (**C**, **G**) two-way ANOVA (**E**, **I**) with Bonferroni post hoc tests. Data presented as ±SEM.

**Fig. S10.**
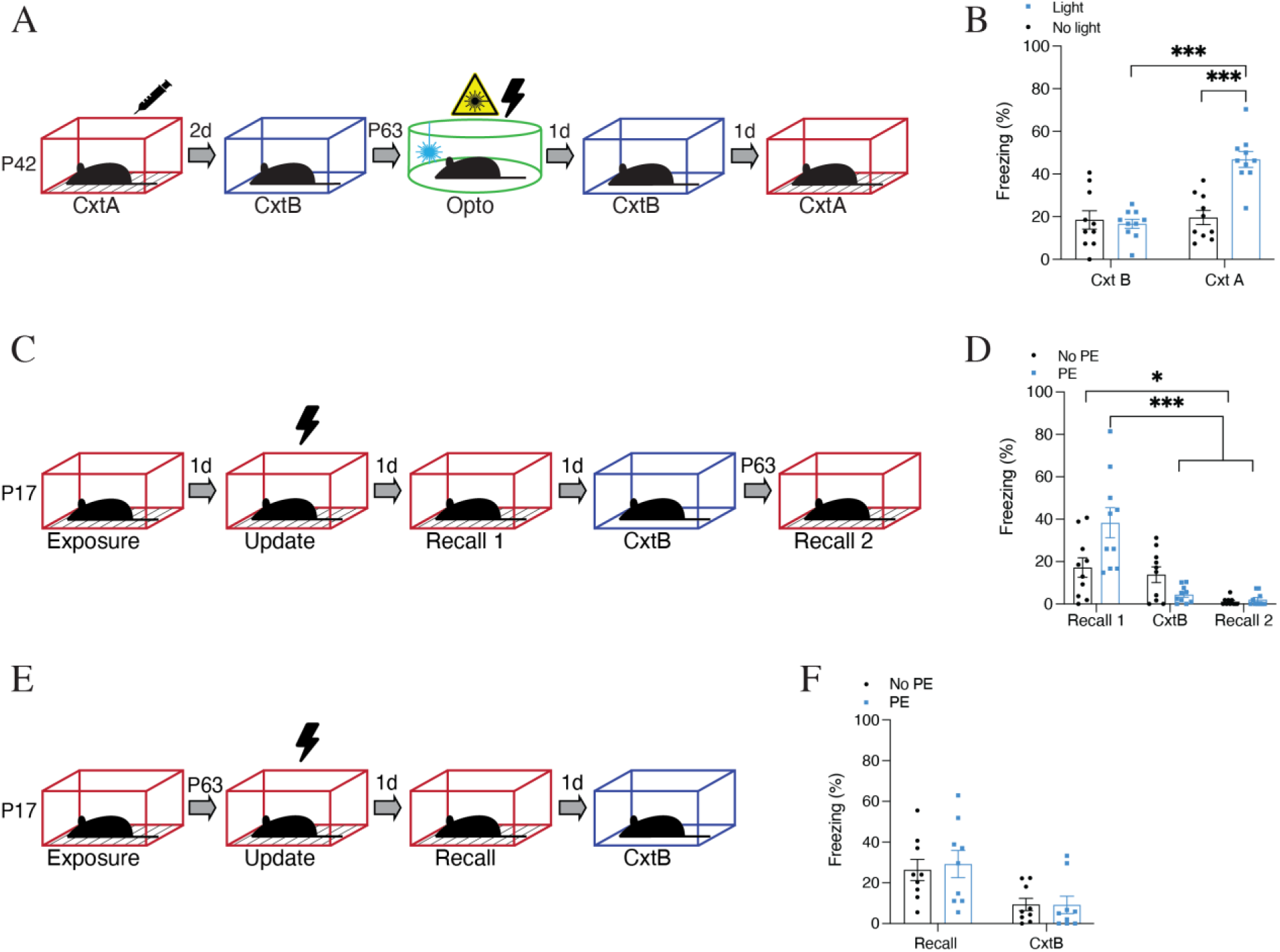
Infant mice can form context-specific memories that can be behaviorally updated. (**A**) Behavioral schedule for artificial updating of an adult engram in Ai32-FosTRAP mice. (**B**) Freezing levels during recall in context B and context A (n=10). Experimental light group froze significantly more in context A (*P < 0.001*). (**C**) Behavioral schedule for updating paradigm in C57BLJ infant mice. (**D**) Freezing levels (n=10) during recall tests at P19 (Recall 1), P20 (CxtB), and P63 (Recall 2). Infant mice that were pre-exposed (PE) to context A froze significantly (*P < 0.05*) more during recall 1. (**E**) Behavioral schedule. (**F**) Freezing levels (n=9) during recall tests. *P < 0.05, P < 0.01, P < 0.001* calculated two-way ANOVA (**B**, **D**, **F**) with Bonferroni post hoc tests. Data presented as ±SEM.

**Fig. S11.**
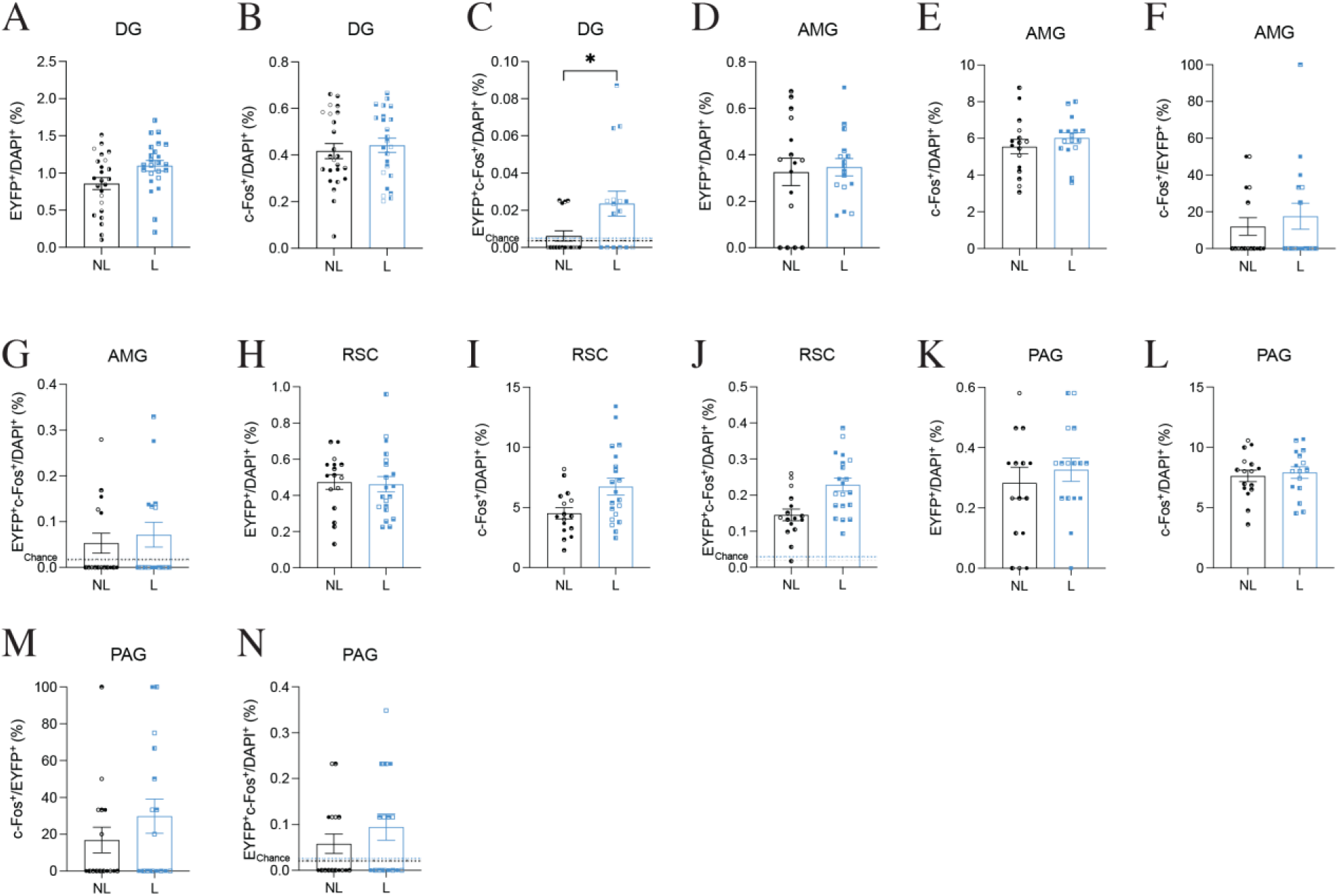
Artificial updating of infant engram cells in adult mice results in an above-chance level of engram reactivation in the DG, AMG, RSC, and PAG. (**A**-**N**) Histological analysis of cell counts after context A recall in adult Ai32-FosTRAP mice. Percentage of (**A**) ChR2-EYFP^+^ and (**B**) c-Fos^+^ cells in the DG (N=6, n=4). (**C**) Percentage of DAPI cells positive for both ChR2-EYFP and c-Fos. The experimental light group showed a significant increase (p < 0.05) in engram reactivation in the DG. Percentage of (**D**) ChR2-EYFP^+^ and (**E**) c-Fos^+^ cells in the AMG (N=4, n=4). Engram reactivation in the AMG as a percentage of (**F**) ChR2-EYFP^+^ and (**G**) DAPI^+^. Percentage of (**H**) ChR2-EYFP^+^ and (**I**) c-Fos^+^ cells in the RSC (N4-5, n=4). (**J**) Percentage is of DAPI cells positive for both ChR2-EYFP and c-Fos. Percentage of (**K**) ChR2-EYFP^+^ and (**L**) c-Fos^+^ cells in the PAG (N=4, n=4). Engram reactivation in the PAG as a percentage of (**M**) ChR2-EYFP^+^ and (**N**) DAPI^+^ cells*. P < 0.05, P < 0.01, P < 0.001* calculated by nested t-test (**A**-**N**). Data presented as ±SEM.

**Fig. S12.**
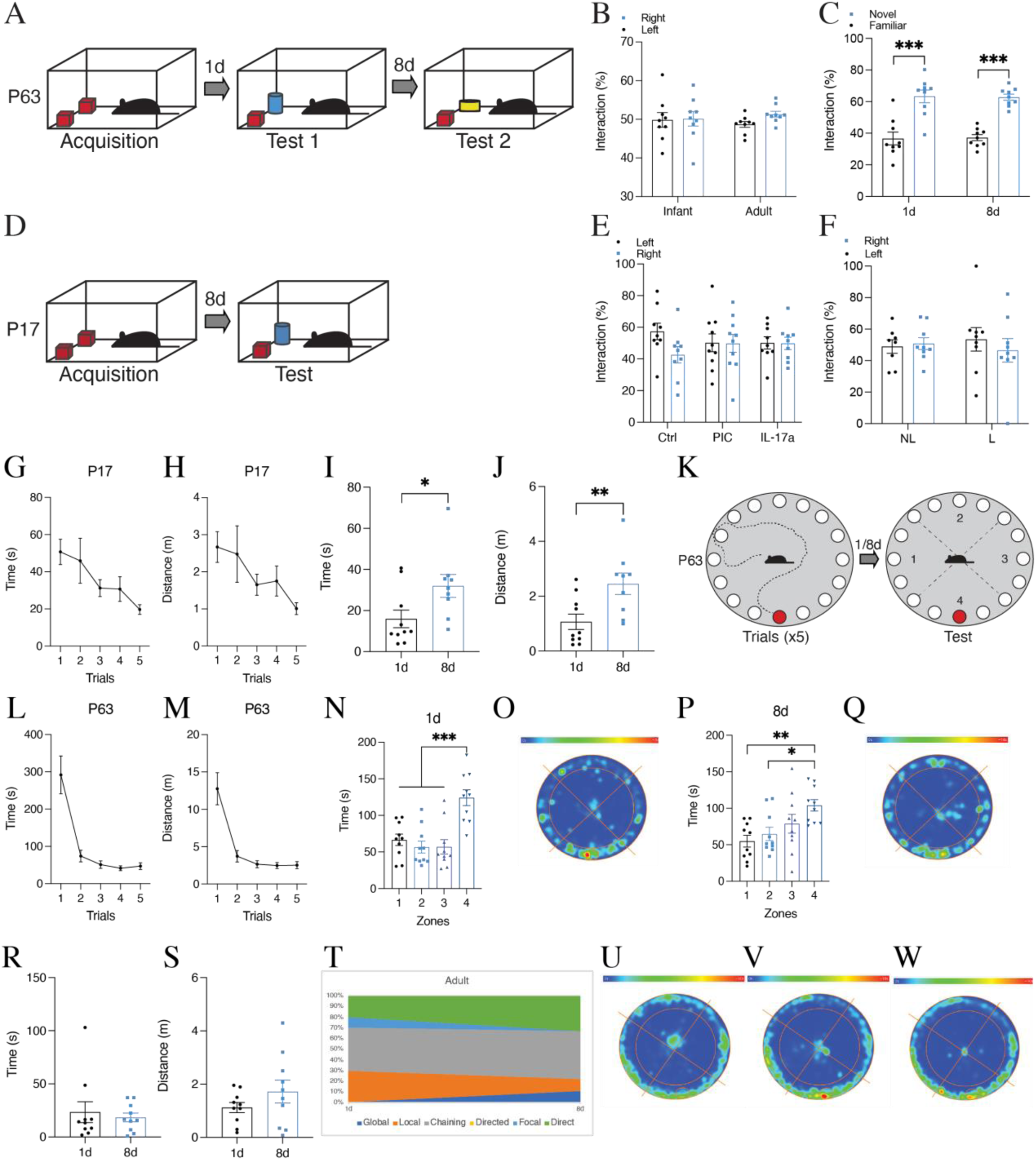
Adult mice retain both an object memory and spatial memory for the location of the escape hole on the Barnes maze task 8 days after training. (**A**) Behavioral schedule for novel object recognition task in adult C57BLJ male mice. (**B**) Object interaction during acquisition in infant (n=10) and adult (n=9) mice. (**C**) Object interaction (%) during novel object recognition test in adult mice (n=9). Mice spend significantly more time exploring the novel object when tested 1 d (p < 0.05) or 8 d (p < 0.001) after acquisition. (**D**) Behavioral schedule for novel object recognition task in infant MIA offspring. (**E**, **F**) Object interaction (%) during acquisition in infant male MIA offspring (n=9-10) and C57BLJ infant mice (8–9). (**G**) Time (s) and (**H**) distance (m) taken by infant C57BLJ mice to reach the escape hole during each trial (n=18-20). (**I**) Time (s) and (**J**) distance (m) taken to first reach where the escape hole should be located during probe tests (n=9-10). (**K**) Behavioral schedule for the Barnes maze in adult C57BLJ male mice. (**L**) Time (s) and (**M**) distance (m) taken to reach the escape hole during each trial (n=20). (**N**-**Q**) Time spent in each zone during probe test (**N**, **O**) 1 d (*P < 0.001*) or (**P**, **Q**) 8 d (*P < 0.05, P < 0.01*) after training. (**R**) Time (s) and (**S**) distance (m) taken to first reach where the escape hole should be located during probe tests. (**T**) Adult male mice search strategies during testing 1 and 8 d after training (n=9-10). (**U**-**W**) Heat map analysis of time spent in each zone by male MIA offspring during probe test 8 d after training (n=9-11) (**U**) Ctrl, (**V**) PIC, (**W**) (IL-17a). *P < 0.05, P < 0.01, P < 0.001* calculated by Student t-test (**I**, **J**, **R**, **S**), one-way ANOVA (**N**, **P**), or two-way ANOVA **(B, C, E, F**) with Bonferroni post hoc tests. Data presented as ±SEM.

**Fig. S13.**
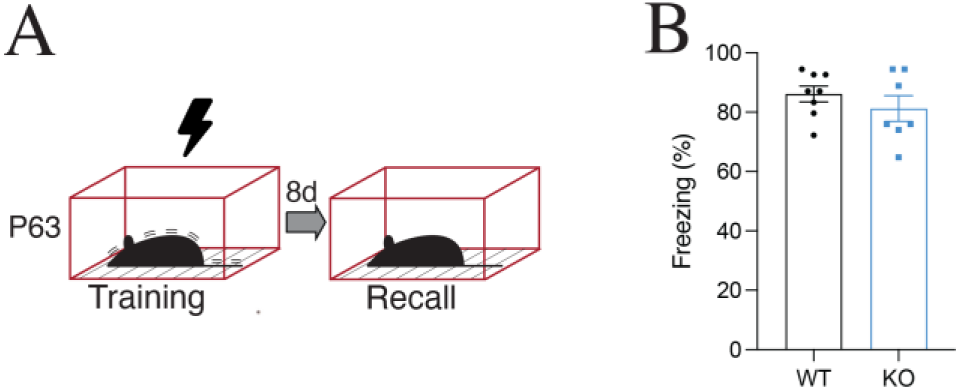
Il17a-/- adult mice show natural memory retention for contextual fear memory when tested 8d after training. (**A**) Behavioral schedule for CFC in Il17a-/- P63 mice. The black lightning symbol represents foot shocks. (**B**) Freezing levels (%) during recall in context A. *P < 0.05, P < 0.01, P < 0.001* calculated by Student t-test (**B**). Data presented as ±SEM.

**Fig. S14.**
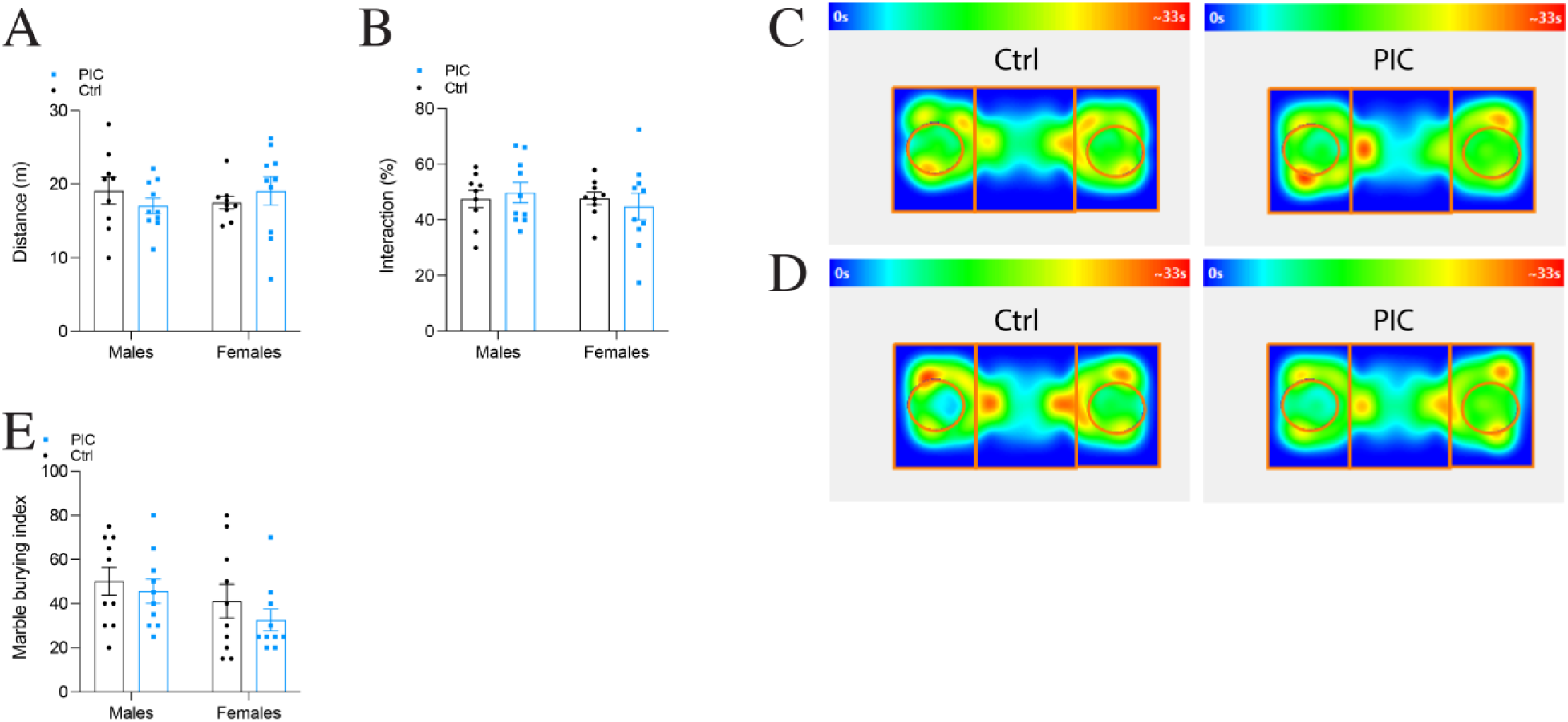
Il17a-/- MIA offspring do not show social interaction deficits or repetitive behavior. (**A**) Total distance (m) traveled during social interaction task in adult male and female (P63) Il17a-/- mice (n=9-10). (**B**) Sociability (%) (**C**) Total distance (m) traveled during social interaction test (n=9-10). (**C**, **D**) Heat map analysis of time spent in each chamber for (**C**) male and (**D**) female Il17a-/- adult MIA offspring. (**E**) Number of marbles buried during marble burying task (n=10). *P < 0.05, P < 0.01, P < 0.001* calculated two-way ANOVA (**A**, **B, E**) with Bonferroni post hoc tests. Data presented as ±SEM.

## References

1. C. Rovee-Collier, The Development of Infant Memory. Curr Dir Psychol Sci. 8, 80–85 (1999).

2. N. S. Newcombe, A. B. Drummey, N. A. Fox, E. Lie, W. Ottinger-Alberts, Remembering Early Childhood: How Much, How, and Why (or Why Not). Curr Dir Psychol Sci. 9, 55–58 (2000).

3. B. A. Campbell, E. H. Campbell, Retention and extinction of learned fear in infant and adult rats. J Comp Physiol Psychol. 55, 1–8 (1962).

4. K. G. Akers, A. Martinez-Canabal, L. Restivo, A. P. Yiu, A. De Cristofaro, H.-L. L. Hsiang, A. L. Wheeler, A. Guskjolen, Y. Niibori, H. Shoji, K. Ohira, B. A. Richards, T. Miyakawa, S. A. Josselyn, P. W. Frankland, Hippocampal neurogenesis regulates forgetting during adulthood and infancy. Science. 344, 598–602 (2014).

5. C. M. Alberini, A. Travaglia, Infantile Amnesia: A Critical Period of Learning to Learn and Remember. J Neurosci. 37, 5783–5795 (2017).

6. B. L. Callaghan, R. Richardson, The effect of adverse rearing environments on persistent memories in young rats: removing the brakes on infant fear memories. Transl Psychiatry. 2, e138 (2012).

7. X. Liu, S. Ramirez, P. T. Pang, C. B. Puryear, A. Govindarajan, K. Deisseroth, S. Tonegawa, Optogenetic stimulation of a hippocampal engram activates fear memory recall. Nature. 484, 381–385 (2012).

8. T. J. Ryan, D. S. Roy, M. Pignatelli, A. Arons, S. Tonegawa, Engram cells retain memory under retrograde amnesia. Science. 348, 1007–1013 (2015).

9. D. S. Roy, A. Arons, T. I. Mitchell, M. Pignatelli, T. J. Ryan, S. Tonegawa, Memory retrieval by activating engram cells in mouse models of early Alzheimer’s disease. Nature. 531, 508–512 (2016).

10. J. N. Perusini, S. A. Cajigas, O. Cohensedgh, S. C. Lim, I. P. Pavlova, Z. R. Donaldson, C. A. Denny, Optogenetic stimulation of dentate gyrus engrams restores memory in Alzheimer’s disease mice. Hippocampus. 27, 1110–1122 (2017).

11. K. Abdou, M. Shehata, K. Choko, H. Nishizono, M. Matsuo, S.-I. Muramatsu, K. Inokuchi, Synapse-specific representation of the identity of overlapping memory engrams. Science. 360, 1227–1231 (2018).

12. T. J. Ryan, P. W. Frankland, Forgetting as a form of adaptive engram cell plasticity. Nat Rev Neurosci. 23, 173–186 (2022).

13. G. B. Choi, Y. S. Yim, H. Wong, S. Kim, H. Kim, S. V. Kim, C. A. Hoeffer, D. R. Littman, J. R. Huh, The maternal interleukin-17a pathway in mice promotes autism-like phenotypes in offspring. Science. 351, 933–939 (2016).

14. C. S. M. Cowan, R. Richardson, Early-life stress leads to sex-dependent changes in pubertal timing in rats that are reversed by a probiotic formulation. Dev Psychobiol. 61, 679–687 (2019).

15. C. S. M. Cowan, A. A. Stylianakis, R. Richardson, Early-life stress, microbiota, and brain development: probiotics reverse the effects of maternal separation on neural circuits underpinning fear expression and extinction in infant rats. Dev Cogn Neurosci. 37, 100627 (2019).

16. S. M. Ohline, W. C. Abraham, Environmental enrichment effects on synaptic and cellular physiology of hippocampal neurons. Neuropharmacology. 145, 3–12 (2019).

17. B. T. Kalish, E. Kim, B. Finander, E. E. Duffy, H. Kim, C. K. Gilman, Y. S. Yim, L. Tong, R. J. Kaufman, E. C. Griffith, G. B. Choi, M. E. Greenberg, J. R. Huh, Maternal immune activation in mice disrupts proteostasis in the fetal brain. Nat Neurosci. 24, 204–213 (2021).

18. R. M. Sullivan, M. Opendak, Defining Immediate Effects of Sensitive Periods on Infant Neurobehavioral Function. Curr Opin Behav Sci. 36, 106–114 (2020).

19. J. H. Kim, R. Richardson, Immediate post-reminder injection of gamma-amino butyric acid (GABA) agonist midazolam attenuates reactivation of forgotten fear in the infant rat. Behav Neurosci. 121, 1328–1332 (2007).

20. J. H. Kim, G. P. McNally, R. Richardson, Recovery of fear memories in rats: role of gamma-amino butyric acid (GABA) in infantile amnesia. Behav Neurosci. 120, 40– 48 (2006).

21. H. H. Y. Tang, G. P. McNally, R. Richardson, The effects of FG7142 on two types of forgetting in 18-day-old rats. Behav Neurosci. 121, 1421–1425 (2007).

22. A. Travaglia, R. Bisaz, E. S. Sweet, R. D. Blitzer, C. M. Alberini, Infantile amnesia reflects a developmental critical period for hippocampal learning. Nat Neurosci. 19, 1225–1233 (2016).

23. A. Guskjolen, J. W. Kenney, J. de la Parra, B.-R. A. Yeung, S. A. Josselyn, P. W. Frankland, Recovery of “Lost” Infant Memories in Mice. Curr Biol. 28, 2283–2290.e3 (2018).

24. R. M. Vlasova, A.-M. Iosif, A. M. Ryan, L. H. Funk, T. Murai, S. Chen, T. A. Lesh, D. J. Rowland, J. Bennett, C. E. Hogrefe, R. J. Maddock, M. J. Gandal, D. H. Geschwind, C. M. Schumann, J. Van de Water, A. K. McAllister, C. S. Carter, M. A. Styner, D. G. Amaral, M. D. Bauman, Maternal Immune Activation during Pregnancy Alters Postnatal Brain Growth and Cognitive Development in Nonhuman Primate Offspring. J Neurosci. 41, 9971–9987 (2021).

25. M. D. Reed, Y. S. Yim, R. D. Wimmer, H. Kim, C. Ryu, G. M. Welch, M. Andina, H. O. King, A. Waisman, M. M. Halassa, J. R. Huh, G. B. Choi, IL-17a promotes sociability in mouse models of neurodevelopmental disorders. Nature. 577, 249–253 (2020).

26. Y. Shin Yim, A. Park, J. Berrios, M. Lafourcade, L. M. Pascual, N. Soares, J. Yeon Kim, S. Kim, H. Kim, A. Waisman, D. R. Littman, I. R. Wickersham, M. T. Harnett, J. R. Huh, G. B. Choi, Reversing behavioral abnormalities in mice exposed to maternal inflammation. Nature. 549, 482–487 (2017).

27. C. J. Guenthner, K. Miyamichi, H. H. Yang, H. C. Heller, L. Luo, Permanent genetic access to transiently active neurons via TRAP: targeted recombination in active populations. Neuron. 78, 773–784 (2013).

28. W. E. Allen, L. A. DeNardo, M. Z. Chen, C. D. Liu, K. M. Loh, L. E. Fenno, C. Ramakrishnan, K. Deisseroth, L. Luo, Thirst-associated preoptic neurons encode an aversive motivational drive. Science. 357, 1149–1155 (2017).

29. L. Madisen, T. A. Zwingman, S. M. Sunkin, S. W. Oh, H. A. Zariwala, H. Gu, L. L. Ng, R. D. Palmiter, M. J. Hawrylycz, A. R. Jones, E. S. Lein, H. Zeng, A robust and high-throughput Cre reporting and characterization system for the whole mouse brain. Nat Neurosci. 13, 133–140 (2010).

30. L. Madisen, T. Mao, H. Koch, J. Zhuo, A. Berenyi, S. Fujisawa, Y.-W. A. Hsu, A. J. Garcia, X. Gu, S. Zanella, J. Kidney, H. Gu, Y. Mao, B. M. Hooks, E. S. Boyden, G. Buzsáki, J. M. Ramirez, A. R. Jones, K. Svoboda, X. Han, E. E. Turner, H. Zeng, A toolbox of Cre-dependent optogenetic transgenic mice for light-induced activation and silencing. Nat Neurosci. 15, 793–802 (2012).

31. J. Li, R. Y. Jiang, K. L. Arendt, Y.-T. Hsu, S. R. Zhai, L. Chen, Defective memory engram reactivation underlies impaired fear memory recall in Fragile X syndrome. Elife. 9, e61882 (2020).

32. R. L. Redondo, J. Kim, A. L. Arons, S. Ramirez, X. Liu, S. Tonegawa, Bidirectional switch of the valence associated with a hippocampal contextual memory engram. Nature. 513, 426–430 (2014).

33. S. Ramirez, X. Liu, P.-A. Lin, J. Suh, M. Pignatelli, R. L. Redondo, T. J. Ryan, S. Tonegawa, Creating a false memory in the hippocampus. Science. 341, 387–391 (2013).

34. M. L. R. and D. A. H. and C. C. Giza, Ontogeny of Rat Recognition Memory measured by the novel object recognition task | EndNote Click, (available at https://click.endnote.com/viewer?doi=10.1002%2Fdev.20402&token=WzM3MTM4NDcsIjEwLjEwMDIvZGV2LjIwNDAyIl0.4_64f4cYvZEqUJwqOd8Yu9gFfDw).

35. B. Bessières, M. Jia, A. Travaglia, C. M. Alberini, Developmental changes in plasticity, synaptic, glia, and connectivity protein levels in rat basolateral amygdala. Learn Mem. 26, 436–448 (2019).

36. A. Garthe, J. Behr, G. Kempermann, Adult-generated hippocampal neurons allow the flexible use of spatially precise learning strategies. PLoS One. 4, e5464 (2009).

37. A. Garthe, I. Roeder, G. Kempermann, Mice in an enriched environment learn more flexibly because of adult hippocampal neurogenesis. Hippocampus. 26, 261–271 (2016).

38. G. Berdugo-Vega, G. Arias-Gil, A. López-Fernández, B. Artegiani, J. M. Wasielewska, C.-C. Lee, M. T. Lippert, G. Kempermann, K. Takagaki, F. Calegari, Increasing neurogenesis refines hippocampal activity rejuvenating navigational learning strategies and contextual memory throughout life. Nat Commun. 11, 135 (2020).

39. M. F. López-Aranda, I. Chattopadhyay, G. M. Boxx, E. R. Fraley, T. K. Silva, M. Zhou, M. Phan, I. Herrera, S. Taloma, R. Mandanas, K. Bach, M. Gandal, D. H. Geschwind, G. Cheng, A. Rzhetsky, S. A. White, A. J. Silva, Postnatal immune activation causes social deficits in a mouse model of tuberous sclerosis: Role of microglia and clinical implications. Sci Adv. 7, eabf2073.

40. T. Dong, J. He, S. Wang, L. Wang, Y. Cheng, Y. Zhong, Inability to activate Rac1-dependent forgetting contributes to behavioral inflexibility in mutants of multiple autism-risk genes. Proc Natl Acad Sci U S A. 113, 7644–7649 (2016).

41. V. Zamoscik, D. Mier, S. N. L. Schmidt, P. Kirsch, Early Memories of Individuals on the Autism Spectrum Assessed Using Online Self-Reports. Front Psychiatry. 7, 79 (2016).

42. B. A. Campbell, J. R. Misanin, B. C. White, L. D. Lytle, Species differences in ontogeny of memory: Indirect support for neural maturation as a determinant of forgetting. Journal of Comparative and Physiological Psychology. 87, 193–202 (1974).

43. L. L. Shook, E. L. Sullivan, J. O. Lo, R. H. Perlis, A. G. Edlow, COVID-19 in pregnancy: implications for fetal brain development. Trends Mol Med. 28, 319–330 (2022).

